# *deCS*: A Tool for Systematic Cell Type Annotations of Single-cell RNA Sequencing Data among Human Tissues

**DOI:** 10.1101/2021.09.19.460993

**Authors:** Guangsheng Pei, Fangfang Yan, Lukas M. Simon, Yulin Dai, Peilin Jia, Zhongming Zhao

**Author notes:** Corresponding authors: (Zhao Z), (Jia P).

## Abstract

Single-cell RNA sequencing (scRNA-seq) is revolutionizing the study of complex and dynamic cellular mechanisms. However, cell-type annotation remains a main challenge as it largely relies on *a priori* knowledge and manual curation, which is cumbersome and less accurate. The increasing number of scRNA-seq data sets, as well as numerous published genetic studies, motivated us to build a comprehensive human cell type reference atlas. Here, we present *deCS* (<u>de</u>coding <u>C</u>ell type-<u>S</u>pecificity), an automatic cell type annotation method augmented by a comprehensive collection of human cell type expression profiles and marker genes. We used *deCS* to annotate scRNA-seq data from various tissue types and systematically evaluated the annotation accuracy under different conditions, including reference panels, sequencing depth and feature selection strategies. Our results demonstrated that expanding the references is critical for improving annotation accuracy. Compared to many existing state-of-the-art annotation tools, *deCS* significantly reduced computation time and increased accuracy. *deCS* can be integrated into the standard scRNA-seq analytical pipeline to enhance cell type annotation. Finally, we demonstrated the broad utility of *deCS* to identify trait-cell type associations in 51 human complex traits, providing deeper insights into the cellular mechanisms of disease pathogenesis. All documents, including source code, user manual, demo data, and tutorials, are freely available at https://github.com/bsml320/deCS.

## Introduction

Recent breakthroughs of single-cell RNA sequencing (scRNA-seq) have greatly boosted our ability to characterize the heterogeneity of cell types in various tissues [1, 2], leading to deeper understanding of disease pathogenesis and development [3, 4]. The typical scRNA-seq data analysis involves unsupervised clustering of the cells, followed by annotation of each cluster [5, 6]. Accurate identification of cell types is critical for downstream analysis. However, this process relies heavily on prior knowledge of cell type marker genes, which is subjective and time-consuming.

In contrast to manual annotation, many automatic annotation tools were recently developed, including *SingleR* [7], *CHETAH* [8], *CellAssign* [9], *scMatch* [10], *scCATCH* [11], *scPred* [12], *scibet* [13], *scVi* [14] and *Cell BLAST* [15]. A recent comprehensive benchmarking evaluation by Abdelaal et al. revealed a large difference in performance among these methods, and such performance was strongly dependent on the reference database [16]. Thus, there is a pressing need to curate high-quality and comprehensive reference panels of human cell type expression profiles, which include the annotated scRNA-seq data from various tissues or sequencing protocols. So far, most of these methods are based on machine learning models, such as support vector machine (SVM) [16] and random forest classifier [17], which are computation-intensive and cannot be directly applied to huge datasets, such as human cell landscape (HCL) [18], human cell atlas [19], and human cell atlas of fetal (HCAF) gene expression [20]. Moreover, the annotation largely relies on a single reference, which may be inaccurate when cell types in “query” and “reference” datasets are not well matched. Therefore, a computational method that can efficiently integrate the annotation results among multiple references is urgently needed. Furthermore, the cell type inference is mostly conducted at single cell level rather than cluster level. The annotation accuracy will significantly decrease, especially for cells with low sequencing depth due to the dropout effect in scRNA-seq data. Even though imputation methods [21, 22] can be applied to recover missing gene expression and improve the annotation accuracy, they are time-consuming [22].

Here, we present *deCS* (<u>de</u>coding <u>C</u>ell type-<u>S</u>pecificity), an automatic scRNA-seq cell type annotation method by decoding cell type-specificity. As an improved version of the *deTS* algorithm [23], *deCS* runs fast, allows the integration of cell annotation from multiple references, and defines cell types at the cell cluster level rather than the single cell level. Specifically, we first created a high-quality and comprehensive human cell type reference panel by including all publicly available large-scale scRNA-seq data sets from various tissues or sequencing protocols, such as HCL and HCAF gene expression. Then, we calculated the *t*-statistic- or z-score-based measurements to “decode cell type-specificity” of gene sets. The *deCS* algorithm was implemented in an R package with different statistical methods. It supports input for either gene expression profiles (e.g., a query scRNA-seq dataset with clusters to be annotated) or list of genes (e.g., a query list of genes). Benchmark results showed that *deCS* reduced computation time and improved annotation accuracy in most tissues compared to other state-of-the-art methods, especially when the cell-type composition of query and reference are not conserved. Therefore, we anticipate *deCS* will become a scRNA-seq routine annotation tool especially when users do not have enough prior knowledge about the cell type markers. Lastly, the curated cell types and their signature genes can be used to explore the cell type-specificity of disease-risk genes. By defining cell type-specific genes (CTGenes) and decoding cell type-specificity, we demonstrated the utility of *deCS* to characterize the relationships between human complex diseases and cell types. Accordingly, these results can provide novel insights into the cellular mechanism of the poorly understood traits and diseases.

## Method

### Data collection for cell type reference panels

#### The BlueprintEncode data

We downloaded the BlueprintEncode RNA-seq data from Aran et al. (2019) [7]. The raw data comprised of 259 bulk RNA-seq samples in total. All cell types were aggregated into 24 broad classes (“main cell types”) with 43 cell types (“fine cell types”). Raw gene expression data was normalized by Transcripts Per Million (TPM), followed by log2 transformation.

#### The Database of Immune Cell Expression data

The DICE reference contained 1561 bulk RNA-seq samples from pure populations of human immune cells. We downloaded the TPM-normalized values for 5 main (15 fine) immune cell types or subtypes from https://dice-database.org/downloads [24].

#### The MonacoImmune data

The MonacoImmune reference comprised 114 PBMCs bulk RNA-seq samples from 4 Singaporean-Chinese individuals. We downloaded the TPM-normalized values for 10 main (29 fine) immune cell types (GSE107011) from Monaco G et al. study [25].

#### The human cell landscape data

The human cell landscape (HCL) reference contained more than 700,000 scRNA-seq expression profiles from more than 50 human different tissues [18]. These cells were grouped into 102 major clusters, and have been well-annotated base on known marker genes. These data were derived from 18 fetal tissues, 35 adult tissues, and several intermediate tissues (e.g., cord blood, placenta). We downloaded the gene expression profiles from http://bis.zju.edu.cn/HCL/landscape.html.

#### The human cell atlas of fetal data

The HCAF gene expression contained ∼4,000,000 single cells from 15 human fetal organs ranging from 72 to 129 days in estimated post-conceptual age [20]. These cells were grouped into 172 major cell types. The raw data is available at https://descartes.brotmanbaty.org/bbi/human-gene-expression-during-development/.

#### CellMatch

The CellMatch reference [11] was curated from various resources, including CellMarker [26], MCA [27], CancerSEA [28], and the CD Marker Handbook. In this study, we only obtained 183 cell types for humans with an average of 163 marker genes per cell type. The raw data set is available from https://github.com/ZJUFanLab/scCATCH/blob/master/R/sysdata.rda.

### Measurement of cell type-specificity of gene

All references derived from bulk RNA-seq were classified into two tiers: tier 1 for the broad (main) cell type category and tier 2 for a more granular (fine) cell type category [7]. For both tiers, we implemented our previous method [23] by fitting a regression model for each gene and computed *t*-statistics to measure the cell type-specificity. Briefly, for cell types in tier 1, we followed our previous work and fitted a regression model for each cell type independently as *Y* ∼ *X*, where *X* is the cell group status (0 or 1) and *Y* is the log2 transformed gene expression. Specifically, for a cell type in the examination, we defined *X* = (*x*_*i*_), *i* = 1,…, *N*, where *N* is the total number of samples; *x*_*i*_ = 1 if the sample belonged to the cell type in the examination, and *x*_*i*_ = 0 if the sample belonged to any cell types that are not in the same group. We then selected the *t*-statistic for the explanatory variable *X* in the standard way:

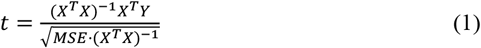

where *MSE* is the mean squared error of the fitted model, i.e.,

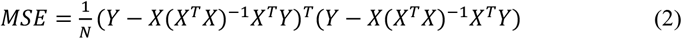

Of note, most cell types in tier 2 were biologically related, such as “*memory B-cells”*, “*naive B-cells”* and “*plasma cells”*. An inclusion of all these instinctively related cell types in one regression model would underestimate their cell type-specificity due to the potential collinearity. Therefore, for the cell types in tier 2, the regression models were fitted after excluding the samples from the same group (main cell type) but keeping the other cell types [23].

The predefined cell number per tissue in the HCL and HCAF data sets is large: 102 (HCL) and 172 (HCAF) major clusters. Accordingly, we utilized *z*-score to measure the cell type-specificity of genes. For each gene, a *z*-score is calculated as *z*_*i*_ = (*e*_*i*_ – mean(*E*))/*sd*(*E*), where *e*_*i*_ is the average log2 transformed expression of the gene in the *i*^th^ cluster, *E* represents the collection of its average expression in all clusters, and *sd* denotes the standard deviation of *E*.

For each cell type, those genes with the highest *t*-statistic or *z*-score are considered as CTGenes in the cell type in examination. The user can define the cutoff values (e.g., the top 5% genes as CTGenes). Because the CellMatch database did not include gene expression information, we directly utilized the predefined CTGenes [11].

To create cell type classification tree, for each reference data set, we conducted cell type hierarchical clustering based on Euclidian distance and ward.D2 linkage aggregation. The bootstrap resampling and probability values (*p*-values) for each cluster was conducted using *Pvclust* package [29].

### Algorithm of cell type-specific enrichment analysis

Feature selection is an important step to determine the cell type. To deal with feature gene collinearity and dropout rate differences, and also to consider variation among datasets, it is better to select features which co-occur in the reference and query data. Although we have already pre-calculated the cell-type specificity score in the reference, we still recommend users to pre-process their query dataset and to use the union of cell cluster-specific genes as input of *deCS*. These recommendations can improve the accuracy and efficiency.

Depending on the query data type, we implemented two test approaches. If the query is an expression profile, we provided two methods to minimize the batch effects between two different datasets, which is a particular concern in scRNA-seq: (1) The z-score strategy normalizes the query expression data by *e*_*n*_ = (*e*_*q*_ - *u*_*s*_)/*sd*_*s*_, where *e*_*q*_ and *e*_*n*_ are the queries and normalized expression, and *u*_*s*_ and *sd*_*s*_ are the mean and standard deviation of a gene from the query expression data. (2) The abundance correction approach [30] normalizes the query RNA-seq data by *e*_*n*_ = *log*_*2*_(*e*_*q*_ + 1)/(*log*_*2*_ (*u*_*s*_ + 1) + 1). After normalization, we calculated the *Pearson correlation coefficient* (PCC) of cell type-specificity of genes between the query and each of the reference cell types, respectively. The most relevant cell type(s), measured by the highest PCC score(s), are annotated to query profiles, possibly with further fine-tuning to resolve closely related cell type(s). On the other hand, we also incorporated a rejection option (e.g., minimum correlation coefficient threshold), which allows the detection of potentially novel cell populations.

When the query is a list of genes (e.g., marker genes of a cell cluster, traits associated genes), we implemented Fisher’s exact test to examine if they are significantly enriched in cell type-specific genes (CTGenes). We allow users to define the threshold, e.g., the top 5% genes ranked by t-statistic or z-score as CTGenes. Specifically, for the query gene set and CTGenes in a given cell type, we build a dichotomous 2 × 2 contingency table as follows:

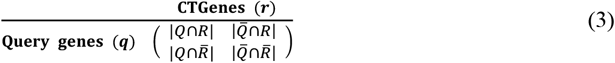

Here, *q* denotes the set of query genes and *r* denotes the set of CTGenes in a given cell type. | *Q*. ∩. *R* | denotes the intersect gene number between *Q* and *R*, 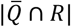 and 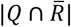 denote the number of genes only in *R* or *Q*, respectively, and 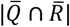 represents the number of genes that neither in *Q* or *R*. In addition, the intersection ratio (IR) |*Q* ∩ *R*| / |*Q*| will also be calculated. An IR of 1 indicates that the 100% query genes are overlapped with CTGenes (default, the top 5% genes) of a given cell type, while 0 indicates no overlap. The most relevant cell type(s), measured by the highest IR(s), are annotated to query profiles.

### Pathway enrichment analysis of cell type-specific genes

For the top 200 CTGenes (genes with the highest *t*-statistic or *z*-score) of each cell type, we used RDAVIDWebService (v 1.19.0) [31] for pathway enrichment analysis. Both Gene Ontology (GO) and Kyoto Encyclopedia of Genes and Genomes (KEGG) pathway annotations are used. Benjamini and Hochberg’s approach [32] was used for multiple test correction. Significant pathways were defined as those with a FDR < 0.05. The networks of CTGenes enriched with KEGG pathways were presented by Cytoscape [33].

### Data collection and preprocessing

#### Single cell RNA-seq data

1. Two PBMC data sets were collected from 10x Genomics website (https://support.10xgenomics.com/single-cell-gene-expression/datasets).
2. The test data set of BIC was collected from healthy controls (HC, n = 3), patients with moderate (M, n = 3) and severe (S, n = 6) COVID-19 infection [34, 35]. The BIC data included 63,103 single cells. The raw data and predefined labels are available from https://github.com/zhangzlab/covid_balf.
3. The FCC scRNA-seq data were collected from 18 human embryos, ranging from 5 weeks (5W) to 25W of gestation [36]. The UMI count of 3842 cells and predefined labels were downloaded from the GEO website (GSE106118).
4. The human liver scRNA-seq data were collected from five fresh hepatic tissues [37]. The raw dataset of 8444 cells and annotation are available through the GEO accession (GSE115469).
5. The scRNA-seq data of human lung, spleen, and esophagus tissues were collected from 12 organ donors [38]. The raw data of 239,224 single cells and annotation are available from https://www.tissuestabilitycellatlas.org/

#### Trait-associated gene lists from GWAS summary statistics

Previous studies have suggested that most trait-associated genes (TAGs) showed strong tissue-specific associations [23, 39]. Therefore, we further collected 51 publicly available genome-wide association studies (GWAS) spanning a wide range of phenotype measurements to investigate the potential association between human traits and cell types.

Considering the statistical noise in GWAS data and more than 90% of genetic variants from GWAS are located in the non-coding regions, for each GWAS trait, we used moderately significant associated SNPs with chi-squared *p*-value < 10^−3^. This strategy allowed us to have a list of genes with weak to strong association signals. Additionally, we applied our DeepFun model [40, 41] to predict their potential regulatory effects. We defined variants with an absolute maximum SAD score > 0.1 as regulatory loci (potential causal variants). Then we employed Pascal software [42] to map them to gene level if these SNPs were located within a range of 50 kb upstream or downstream of corresponding gene transcription start sites by taking into account linkage disequilibrium (LD) and gene length information. Any genes with at least one regulatory locus were regarded as TAGs.

#### Bulk RNA-seq data

The test bulk RNA-seq data were generated from schizophrenia associated human iPSC-derived cell lines from the population isolate of the Central Valley of Costa Rica [43]. One clone from each subject was differentiated into NPCs and neurons, respectively. In total, we collected RNA-seq data from iPSC-NPCs (n = 13) and iPSC-neurons (n = 11).

### Single cell permutation analysis

Single-cell permutation analysis was performed to assess the cell cluster internal correlation and detected gene richness. For a given number (n from 1 to 50) of cells belonging to the same cell type, we performed random sampling 100 times without replacement, then calculated the PCC of the averaged gene expression level from the single cells compared to the pseudo-bulk level (cell cluster averaged). In addition, we estimated the total number of detected genes that could be identified (at least one UMI in at least one cell), along with the cumulated number of cells.

### Statistical analysis

UMAP analysis [44] was used to visualize scRNA-seq batch effect. For comparison of the cell annotation accuracy among healthy controls, moderate, and severe COVID-19 infected patients, we used *kruskal*.*test* function in *R* software. The performance of *deCS* was evaluated by recall, defined as TP/(TP + FN) where TP and FN denote the number of true positives and false negatives.

### Evaluation and software implementation

The evaluation was conducted on a desktop equipped with i7-7700HQ CPU and 16 GB of RAM. We ran *deCS* using two approaches: correlation analysis and Fisher’s exact test. *deCS* runs fast when applying correlation analysis. It took only ∼7 seconds for a gene expression matrix with BIC data set (63,103 cells) by *deCS*.*correlation* function. When applying Fisher’s exact test, it took only ∼50 seconds for a list of 43,514 genes across the 51 traits.

## Results

### Overview of *deCS* workflow

Our goal is to develop a tool to perform automatic cell type annotation across datasets of different sequencing protocols and levels of complexity. To this end, we first collected various public cell type expression profiles. As shown in **Figure 1**, we included several public human bulk RNA-seq data such as BlueprintEncode [45, 46], the Database of Immune Cell Expression (DICE) [24], and MonacoImmune [25]. Although these were bulk data, they were generated using cell lines and have already been used as reference datasets in *SingleR* [7]. In addition, we downloaded the most comprehensive human single-cell expression data resource from the human cell landscape (HCL) [18] and human cell atlas of fetal (HCAF) projects [20]. For each data set, we applied different analytical strategies (see Materials and Methods for more details) to define CTGenes. Finally, we integrated the manually curated marker genes (used as CTGenes in *deCS*) from the CellMatch database [11]. Based on genes *t*-statistic/*z*-score and CTGenes, we implemented two statistical tests for cell type annotation (Figure 1). If the query data is a “gene expression profile” with unknown cell types, *deCS* applied the correlation analysis to determine which cell types in the reference are significantly similar to those in the query data. If the query data is a set of “candidate genes” of interest, *deCS* determines if they overlap significantly with a cell type in the reference by using Fisher’s exact test.

**Figure 1.**
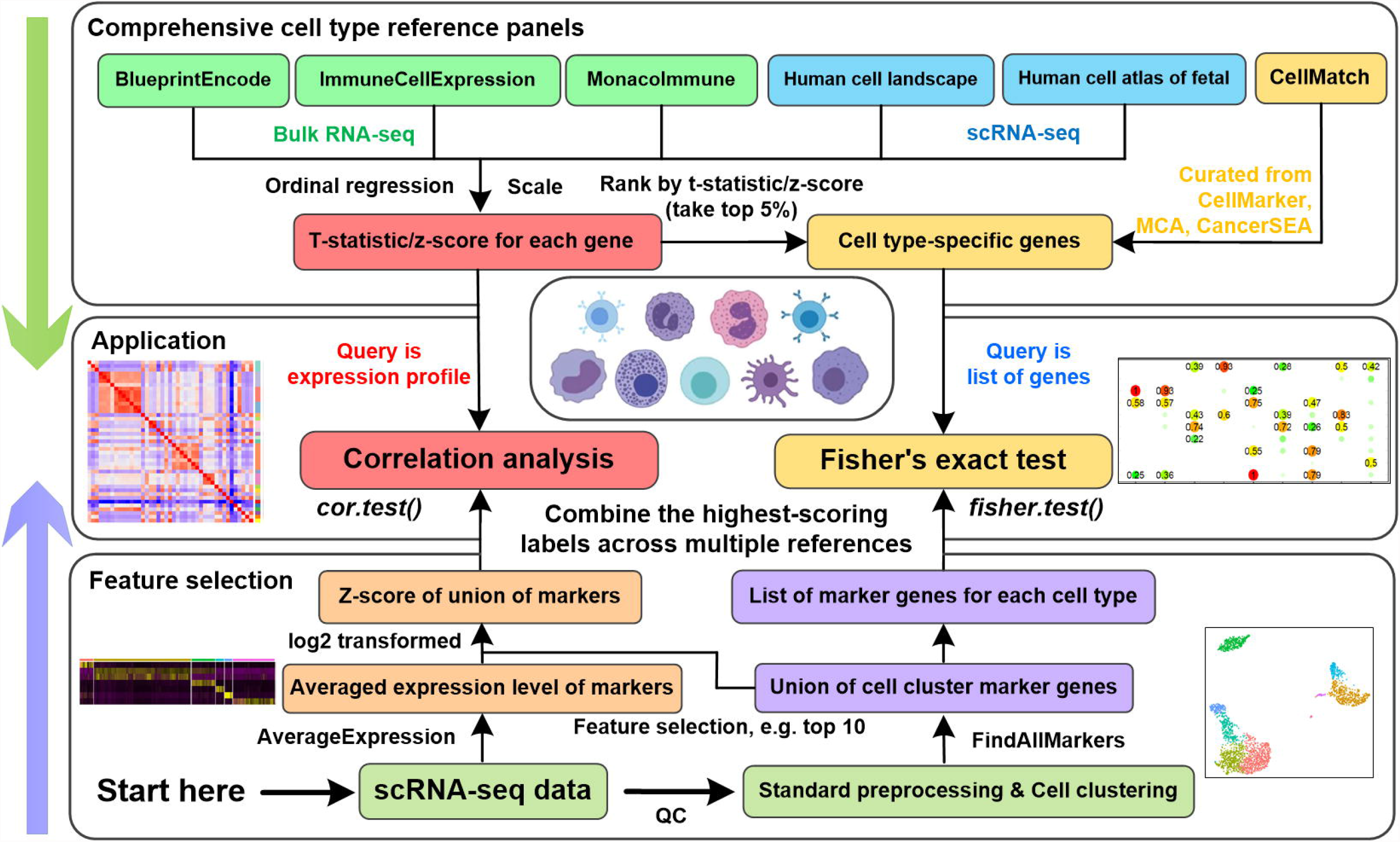
Overview of *deCS* flowchart. For each cell type, we compute *t*-statistics and *z*-scores for each gene in the bulk RNA-seq and scRNA-seq derived references, respectively. Then, we define genes with the highest *t*-statistics or *z*-scores (top 5%) as CTGenes, cell type-specific genes. We further integrate cell signature gene sets from one curated CellMatch database. Depending on the type of “query data”, when the query input is gene expression profile, it will calculate the *Pearson* or *Spearman’s rank correlation coefficient* between query scaled expression profiles and *t*-statistics (or z-scores) of each cell type in the reference, then assign the label with the highest score to the query profile. When the query input is list of genes, it is analogous to existing tools for identifying candidate genes that are overrepresented in specific GO terms or KEGG pathways [31]. Finally, the top enriched cell type is annotated to query data.

### Cross-reference validation and biological pathway enrichment of CTGenes

To validate the reliability and consistency of CTGenes, we first calculated internal cell type correlation and then utilized Uniform Manifold Approximation and Projection (UMAP) dimension reduction analysis [44] to visualize the global landscape of expression profiles (Figure S1). We further validated the consistency of CTGenes across different references, including BlueprintEncode, DICE, MonacoImmune, and HCL. Accordingly, we systematically compared the CTGenes (by default using the top 5% highest genes by *t*-statistic) between two different references. We observed a large variation between references, with matching rate ranging from 20% to 70%, although more than 80% of matched cell types shared the highest cell type-specific marker genes (Figure S2). For example, when comparing the BlueprintEncode reference with the HCL reference, only 36 out of 102 (35.3%) cell types in HCL shared at least 200 CTGenes with the BlueprintEncode reference, while most cell types in HCL reference [e.g., alveolar type 2 (AT2) cells in lung, mast cells in blood] were missed in BlueprintEncode.

To further assess the validity of CTGenes identified by *t*-statistic, we performed pathway enrichment analysis for each cell type using the top 200 CTGenes (Table S1). We constructed a network containing all significantly enriched pathways (false discovery rate (FDR) < 0.05) and corresponding cell types. As shown in **Figure 2**, more than 90% CTGenes were correctly enriched in biologically relevant pathways: adipocytes were enriched in “*Regulation of lipolysis in adipocytes*”; B-cells were enriched in “*B-cell receptor signaling pathway*”; CD4+ and CD8+ T-cells were enriched in “*T-cell receptor signaling pathway*”; HSCs were enriched in “*Spliceosome*”; melanocytes were enriched in “*Melanogenesis*”; and neuron cells were enriched in “*GABAergic synapse*” and “*Glutamatergic synapse*”. In addition, the relationship of CTGenes could be observed based on a force-directed layout network. For example, several cell types were well clustered: myocytes and skeletal muscles; CD4+, CD8+ T-cells and NK cells; and monocytes, dendritic cells (DC), and macrophages.

**Figure 2.**
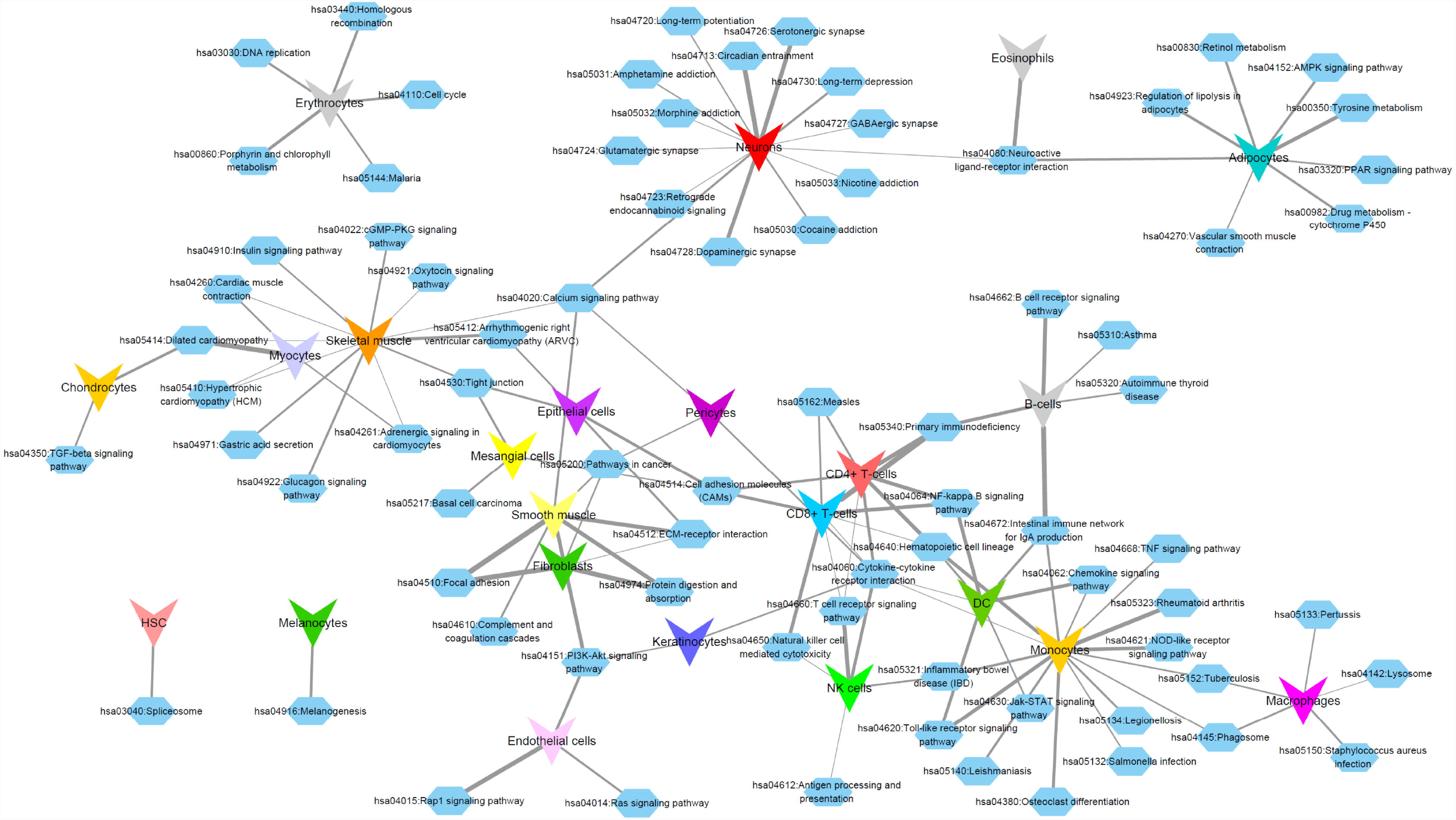
KEGG pathway enrichment of cell type-specific genes. The network shows the relationship between CTGenes and KEGG pathways with significant association (FDR < 0.05). The network layout is based on a force-directed graph. Edge width is proportional to -log_10_(transformed FDR).

### scRNA-seq cell type annotation

#### Peripheral blood mononuclear cells

Human peripheral blood mononuclear cells (PBMCs) are routinely studied in the biomedical fields including immunology and the development of diagnostics and therapeutics for human diseases [47]. To explore the utility of *deCS* for direct annotation of scRNA-seq data, we first analyzed a PBMC data set, which is available from 10x Genomics using the MonacoImmune reference. The basic workflow of two alternative annotation pipelines is depicted in Figure S3. We followed the standard pre-processing workflow for PBMC scRNA-seq data in Seurat [5]. We extracted the union of top 10 cell cluster-specific marker genes for each cluster, and then calculated the average expression profile followed by z-score normalization (Figure 1). We applied two statistical approaches: correlation analysis and Fisher’s exact test would match the cell label with the highest PCC or intersection ratio (IR) to each query cluster (**Figure 3**). Overall, the main labels predicted by all 9 clusters were the same between these two methods on the PBMCs data set. The similarity scores of most clusters mapped to a single main label and were significantly higher compared to the remaining cell types, indicating high specificity on main cell label annotation. When mapping to fine labels, many cell clusters were mapped to multiple fine labels. Nevertheless, most of them were mapped to closely related cell types. For example, native CD4 T-cells were predicted as one subtype CD4 T-helper 1 cells (Th1) [48].

**Figure 3.**
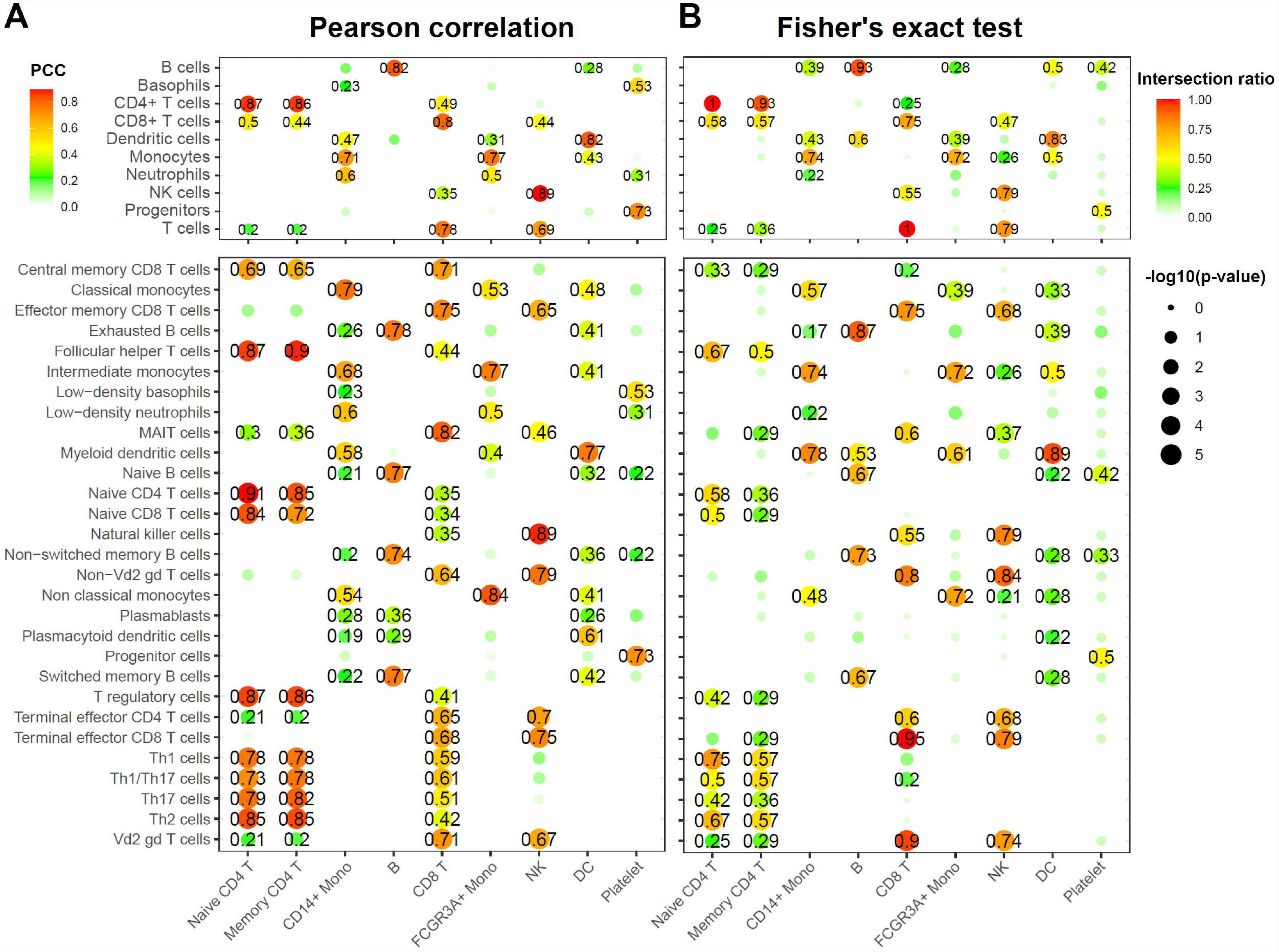
Comparison of the two methods in *deCS* using PBMC scRNA-seq data. **A**. Correlation analysis by PCC. **B**. Fisher’s exact test. *X*-axis: 9 cell type clusters (query). *Y*-axis: the 10 main (up panel) and 29 fine (bottom panel) immune cell types from the MonacoImmune reference [25]. The colors represent the PCC or genes intersection (IR) ratio, and the sizes represent the -log_10_(transformed *p*-value). Non-significant associations (*p*-value > 0.05) were labeled by white color.

#### Solid tissues on adult and fetal samples

We next evaluated the performance of *deCS* in six additional scRNA-seq data sets from different solid tissues. We first considered bronchoalveolar immune cells (BIC) from COVID-19 infected patients [34]. The results indicated that *deCS* found the best-matched cell type for the majority of cell clusters (8 out of 10) on BlueprintEncode panel (**Figure 4**A, B). Two exceptions were mast cells and plasmacytoid dendritic cells (pDC), which were matched to erythrocytes (PCC = 0.5, IR = 0.25) and B cells (PCC = 0.28, IR = 0.2), respectively. Here, IR refers to the intersection ratio. One major concern of the misclassification is that cell type is not represented in the used BlueprintEncode reference [8]. One possible solution is to annotate them as undetermined cells if they are too dissimilar to any references (e.g., PCC < 0.3) [49]. Alternatively, users can map their query data to other available references to identify novel cell types. As shown in Figure S4, mast cells and pDC were mapped to their counterpart when using the HCL (PCC = 0.82, IR = 0.77) and MonacoImmune (PCC = 0.74, IR = 0.9) references, respectively.

**Figure 4.**
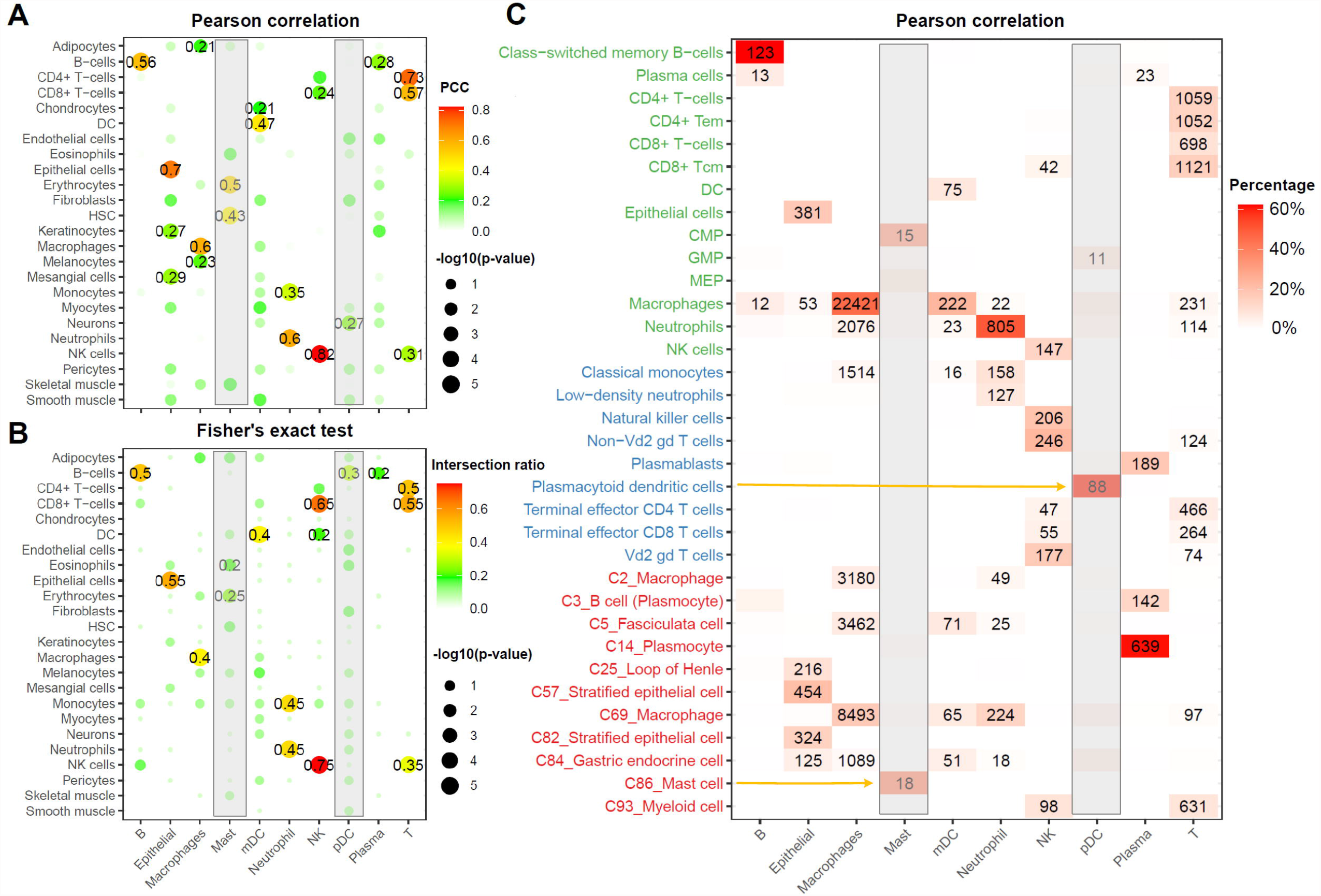
*deCS* annotation of bronchoalveolar immune cells from COVID-19 infected patients. **A**. Correlation analysis by PCC. **B**. Fisher’s exact test between 10 cell major clusters marker genes and CTGenes in BlueprintEncode reference. *X-*axis: 10 cell type clusters (query). *Y*-axis: the 24 main cell types from BlueprintEncode. The colors represent the PCC or genes intersection ratio, and the sizes represent the -log_10_(transformed *p*-value). **C**. Individual cell level annotation of 63,103 single cells profiled on the 10x Genomics platform. We annotated each cell by using the top matched mode (cell type with the highest PCC). Using integrated references (BlueprintEncode: green; MonacoImmune: blue; HCL: red), *deCS* classifies all cells that are nearly identical to that by manual classification. The colors and the numbers representing the percentages and the total cell counts were annotated to a given cell type.

The second data set was derived from fetal cardiac cells (FCC) [36]. As shown in Figure S5, almost half cell clusters (5/9) failed to find any matched cell types (using PCC > 0.3 as the threshold) when mapping to BlueprintEncode. However, when using the HCL or HCAF reference, there were 8 out of 9 cell clusters (except for C5 valval cell) mapped to the annotated cell type. Similarly, for the four human scRNA-seq datasets of liver, lung, spleen, and esophagus tissues, only 50∼70% of cell clusters were mapped to their counterpart when using BlueprintEncode, HCL or HCAF reference individually (Table S2). When combining the highest-scoring labels across multiple references, mapping rate was improved to 87.8% or above. This demonstrated that integrating results from multiple high-quality curated reference panels could effectively identify novel cell types that cannot be detected by other methods, thereby significantly improve annotation accuracy and utility.

### Annotation in single cell resolution

In the above analysis, we used the averaged expression level in each cluster to evaluate the performance of *deCS* [50]. These traditional routines, however, overlook an important characteristic of scRNA-seq data [51]: cellular heterogeneity. Considering the true hierarchical cluster structure for a cell population (e.g., both CD4+ and CD8+ are T cells) [51], cellular heterogeneity is widely observed when applying unsupervised clustering [52]. The ideal scenario is each cell can be annotated to one specific branch of the hierarchical tree [8]. Therefore, we further evaluated *deCS* in BIC data set [34]. As shown in Figure 4C, the majority of cells within 10 clusters can be matched to the best cell type through the integrative analyses across multiple references. There may exist misclassifications only in closely related cells. For example, for myeloid cells, 1514 and 2076 cells in macrophages cluster were erroneously annotated as monocytes and neutrophils, respectively. Some neutrophils were annotated as monocytes or macrophages (Figures S6, S7).

Previously the scMatch method [10] showed that the cells with more reads were more likely to be correctly classified than those with lower read depth. Therefore, based on the sequencing depth, we divided 63,103 cells from BIC data into three groups: low (1k ≤ UMI < 3k), median (3k ≤ UMI < 6k), and high (UMI ≥ 6k) depth groups. As **Figure 5**A showed, although low sequencing depth has a slight impact on annotation accuracy (e.g., 10 B cells were annotated to macrophages), *deCS* approach had a good performance for other cell types. As a negative control, we further compared the annotation recall among healthy controls and moderate or severe COVID-19 infected patients. As shown in Figure 5B, the cell type frequencies among different groups demonstrated significant differences [Kruskal–Wallis test (p < 0.05)], especially for epithelial and neutrophil cell types [53].

**Figure 5.**
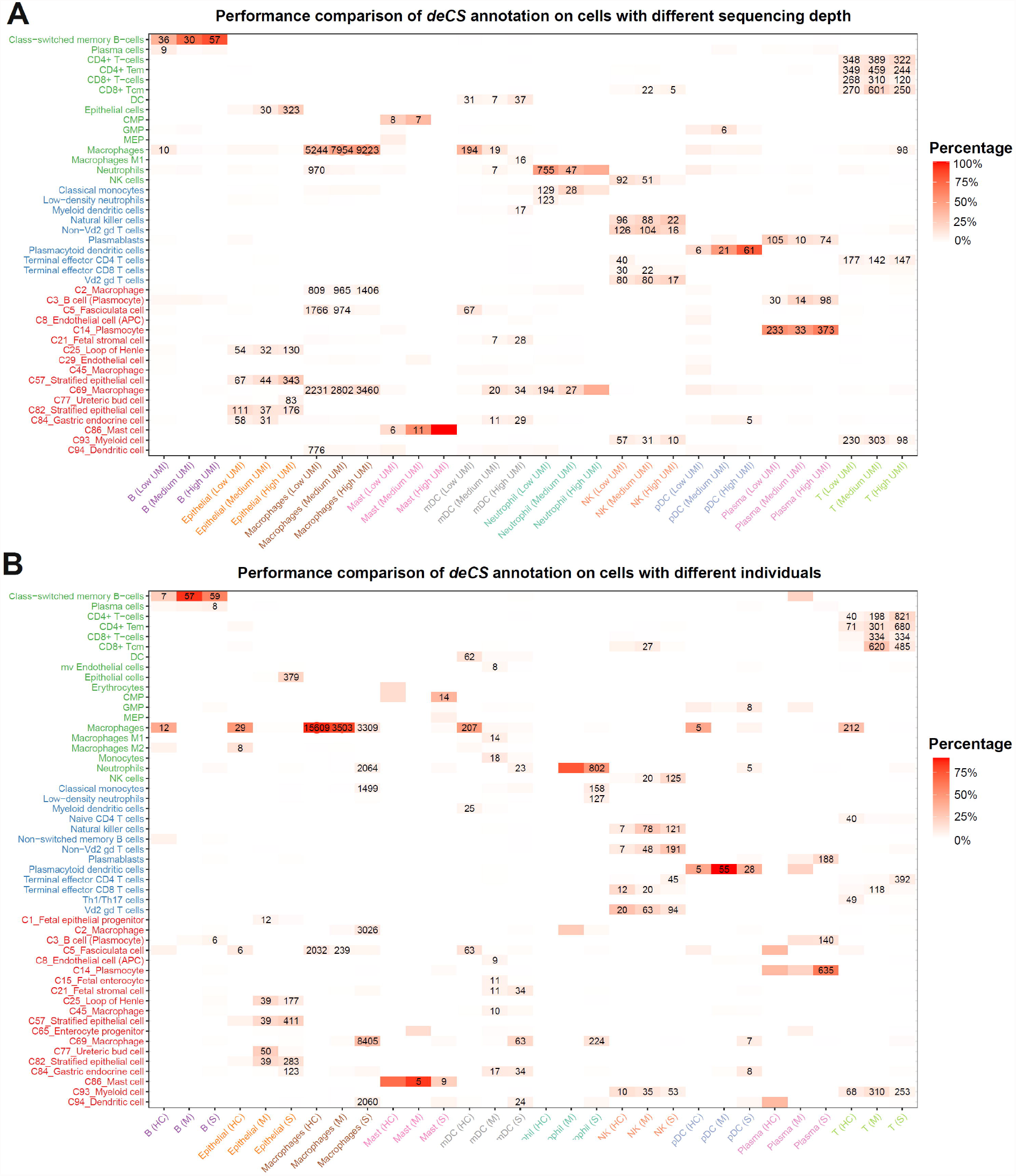
Comparison of *deCS* annotation on cells with different sequencing depth and individuals. **A**. Annotation comparison of cells sequenced among low (1k ≤ UMI < 3k), median (3k ≤ UMI < 6k), and high (UMI ≥ 6k) depth groups. **B**. Annotation comparison of cells derived from HC (healthy controls), M (moderate), and S (severe) COVID-19 infection groups. We only use the union of top 20 marker genes of each cluster. The colors and the numbers represent the percentages and the total cell counts that were annotated to a given cell type. Only cell types are annotated with at least 5% cells from 10 predefined cell clusters were labeled with the number of cell counts.

Although *deCS* demonstrated good performance at the single-cell level, there remained a small fraction of misclassified cells. To address this issue, we further investigated top marker genes for each cluster to better understand the underlying reason. As shown in Figure S8, the top marker gene of naive CD4+ T cell, *CCR7* was expressed in approximately 40% of the naive CD4+ T cells due to the dropout effect in scRNA-seq. The annotation accuracy was decreased when cell type inference was conducted at single cell level. While utilizing gene expression imputation approaches [21, 54, 55] could improve single cell resolution annotation performance. Thus, we recommend cell type annotation on clusters instead of individual cell after imputation, which is consistent with other publications [5, 56].

### Benchmarking analysis on different tissues

To demonstrate the superior performance of *deCS*, we compared it with other methods, including *SingleR* [7], *CHETAH* [8], *scPred* [12], *scibet* [13], *scVi* [14] and *Cell BLAST* [15] using the above eight datasets. We first applied the methods to two PBMC data sets. As summarized in **Table 1**, the overall performances of *deCS, SingleR, scPred* and *sciBet* ranged from 87.55% to 89.41%. Among them, *scPred* was the best-performing classifier (88.78%), which is consistent with previous benchmark work, demonstrating that SVM classifiers have overall the best performance [16]. However, as classical genes are not expressed in every single cell, *deCS* showed better performance (89.41%) than *scPred* (Table S3) after imputation [21]. In addition, we noticed that *CHETAH, Cell BLAST*, and *scSorter* had weaker robustness on the annotation accuracy. One possible reason is that the rejection threshold is too strict, such that most cells were annotated to be ambiguous or an intermediate branch of cell types. Another possible reason is potential overcorrection when some query cell types are non-overlapping with reference (details are shown in Table S2 and Figure S9).

**Table 1.** Performance comparison of *deCS* with six other models among eight data sets.

| ID | Tissue name | deCS(Cor) | deCS(FET)* | SingleR | scPred | sciBet | CHEATH | Cell blast | scSorter** |
| --- | --- | --- | --- | --- | --- | --- | --- | --- | --- |
| 1 | PBMC1 | 88.74% | - | 87.81% | 88.78% | 87.55% | 54.34% | 77.38% | 75.81% |
| 2 | PBMC2 | 88.52% | - | 87.81% | 88.59% | 87.79% | 54.42% | 78.40% | 71.86% |
| 3 | Bronchoalveolar | 97.34% | 95.79% | 71.34% | <50% | 91.54% | <50% | <50% | - |
| 4 | Heart | 80.97% | 80.97% | 63.33% | <50% | <50% | <50% | <50% | - |
| 5 | Liver | 99.56% | 92.37% | <50% | <50% | 83.99% | <50% | <50% | - |
| 6 | Lung | 98.59% | 87.78% | 70.72% | <50% | 67.77% | <50% | <50% | - |
| 7 | Oesophagus | 87.64% | 87.07% | 86.82% | <50% | 96.34% | <50% | <50% | - |
| 8 | Spleen | 97.34% | 90.54% | 91.90% | 79.63% | 90.35% | <50% | <50% | - |

*Note:* \* We benchmarked PBMC data sets at the single cell level, because *deCS*(FET) model needs cell-cluster level information, which is not available from these PBMC data sets. \*\* scSorter is a “semi-supervised” cell type assignment method. Because the prior knowledge of markers would strongly affect the annotations, we did not benchmark scSorter on six solid tissues.

Since most methods showed comparable performance on PBMC, we further benchmarked them on six extra datasets, including bronchoalveolar, heart, liver, lung, oesophagus, and spleen. As described in **Table 1**, the *deCS* correlation analysis showed an average of 93.6% accuracy. When *deCS* Fisher’s exact test failed to consider the “weight” feature, it only obtained an average of 89.1% accuracy. *sciBet* (86.0%) *and SingleR* (76.8%) also achieved good performance on most tissues, as both of them incorporated a comprehensive reference panel with middle size (>30 cell types). However, the accuracy for most other methods was lower than 50% due to inappropriate reference panels, even when counting those undetermined cells (when query cell types are non-overlapping with reference). For example, as most cell types from spleen were identical with PBMC, *scPred* showed 79.6% accuracy on spleen tissue. However, on liver tissue, where there was a large proportion of hepatocyte (48.2%) (non-overlapping with reference), the accuracy was below 50%, as 41.3% of the hepatocytes were annotated to monocytes (rather than unassigned cell type). In most studies, researchers typically do not have enough prior knowledge about the cell type composition in the investigated samples. Therefore, the expanded reference in *deCS* can be widely used on various tissues.

### *deCS* reduced the feature collinearity and impact of different sequencing depth

The superiority of *deCS* is not only due to the expanded cell type. Even after excluding those query-specific cell types (e.g., ventricle cardiomyocytes in heart, hepatocytes in liver, and alveolar cells in lung), the *deCS* showed better annotation accuracy in distinguishing biologically related cell types (e.g. CD4+, CD8+ T-cells and NK, monocyte, and macrophage). We speculated feature selection (comparing to *SingleR*) is still an essential step before correlation analysis. These identified anchor genes (cluster marker genes) can effectively reduce the collinearity compared to the case when using more genes (top variable genes). As shown in **Figure 6A**, the query cluster is naive CD4+ T cells and the PCC was 0.87 when using “union of cluster-specific genes”, but dropped to 0.45 when using top 2000 variable genes. In contrast, the PCC between query cluster and CD8+ T cells was still high (0.32) (Figure 6B). Although there are lots of other methods for feature selection and model prediction (e.g., SVM) [12, 13], it is not realistic to conduct such time-consuming process on huge HCL or HCAF raw data (> 4 million cells) [20]. In consideration of both speed and scalability when combining the highest-scoring labels across multiple references, we decided to provide the cell cluster aggregated *t*-statistic and *z*-scores. On the other hand, *deCS* drastically reduced the required memory and runtime, which is helpful for annotation of large data set using a personal computer.

**Figure 6.**
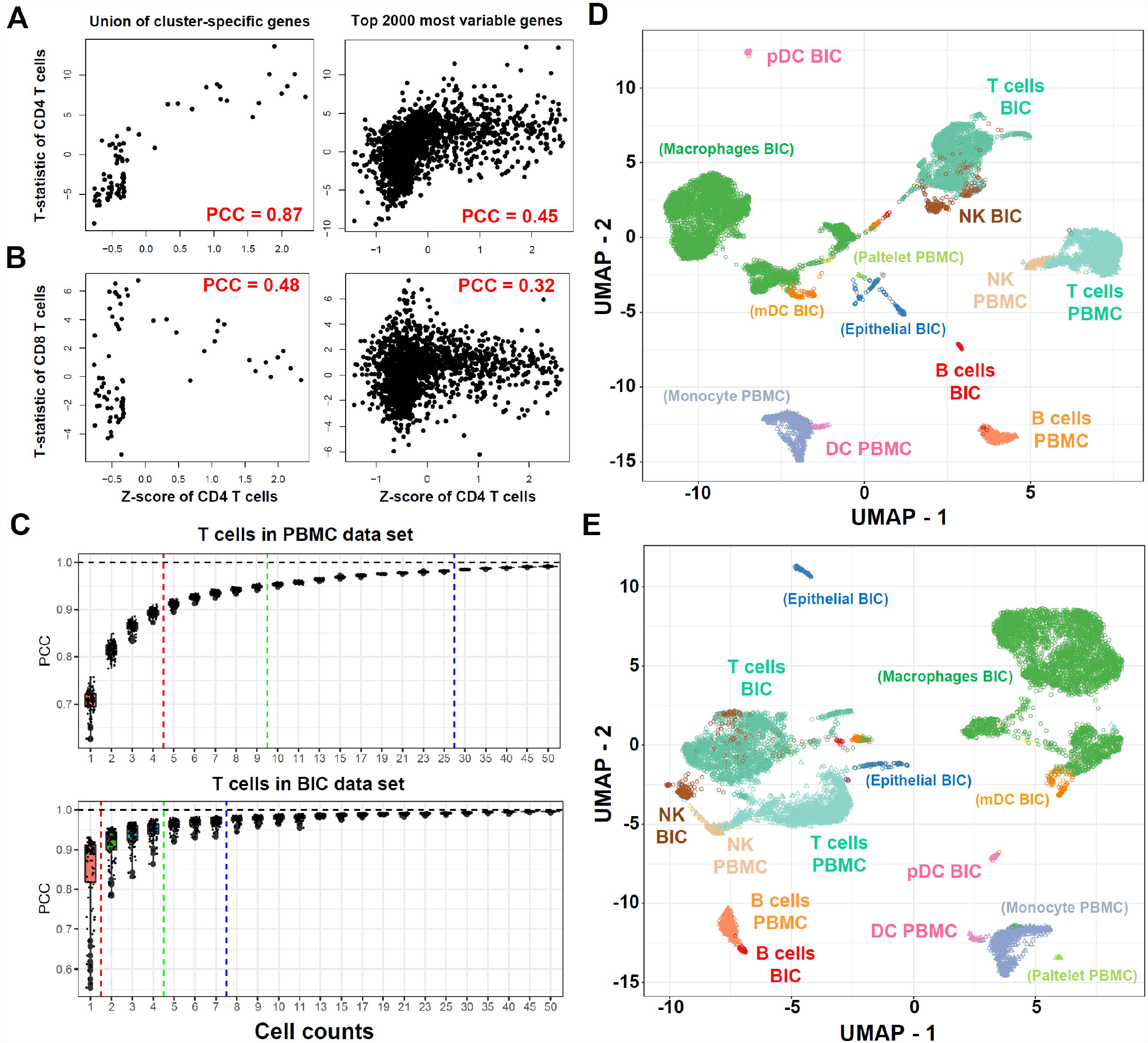
*deCS* pipeline reduces the collinearity and impact of sequencing depth difference. **A**. Correlation comparison by using the union of cell cluster-specific gene and top 2000 most variable genes between query (CD4+ T) cells and reference on CD4+ T (positive) or **B**. CD8+ T cells (negative). **C**. Rarefaction curve of T cells in PBMC and BIC data sets. *X-*axis: the sized pools (n from 1 to 50). *Y*-axis: the inter-correlation between the gene expression of pooled cells and bulk level (cluster averaged). Plotted are IQR, interquartile ranges. The red, green, and blue lines indicate the number of single cells that hypothetically enabled the averaged expression levels of pooled cells are approximate to bulk populations (with PCC > 0.90, 0.95, and 0.975). **D**. UMAP, Uniform manifold approximation and projection integrative analysis of PBMC and BIC data sets by log normalized expression matrix or **E**. further z-score scaled. Each circle and triangle represents a single cell derived from BIC or PBMC data sets.

Due to the stochasticity and inefficient mRNA capture in scRNA-seq, varying sequencing depth across batches is a major driver of batch effects [7, 57]. Thus, we evaluated the level of sequencing depth (different sized pools, n from 1 to 50) of single cell transcriptomes to the pseudobulk level (cluster averaged). As shown in Figure 6C, we observed a gradually diminishing improvement of inter-correlation with increasing sequencing depth. For example, the inter-correlation between single cell and pseudobulk level for T cell in PBMC and BIC data sets was 0.707 and 0.898, respectively. As the median number of detected genes in BIC was almost 4-fold more than PBMC data set, it only needed to cumulate 5 cells to approximate 95% bulk level in BIC, while it needed at least cumulated 10 cells in PBMC data (Figure 6C). To put it another way, when a gene is observed at a low or moderate expression level in BIC cells, it is probably not detected in PBMC cells. This phenomenon is also known as “dropout” [58]. For better demonstration, we randomly selected 5 bulk RNA-seq samples and down-sampled each sample to 10-100k reads followed by quantification normalization (Table S4). Interestingly, in the low sequencing depth condition, we noticed a pronounced effect of “sequencing depth” compared to the “identity” level in the PCA plot. In contrast, z-score normalization effectively reduced the “batch effect” (Figure S10). Furthermore, standard UMAP analysis was applied to two different data batches (PBMC and BIC). As shown in Figure 6D, we observed a strong batch effect between PBMC and BIC log normalized expression matrices (e.g., NK and T cells were clustered by two samples). In contrast, after z-score normalization (Figure 6E), most cell types form distinct clusters are separated by cell types (e.g., B cells, NK cells, T cells, and DC cells).

### Decoding the cell type-specificity on other applications

In addition to scRNA-seq cell type annotation, *deCS* reference panels can be used to detect cell-type specificity and to provide insightful understanding of many other biological problems. For example, we can apply *deCS* to genome-wide association studies (GWAS) data for trait-cell type associations, and also bulk RNA-seq data from induced pluripotent stem cell (iPSC) derived cells. We demonstrated the utility of *deCS* with two applications below.

Understanding the underlying context of human complex diseases is an important step to unveil the etiology of disease origin [59], yet tissues are complex milieus consisting of various cell types [60]. Therefore, tissue level association may fail to elucidate cell type contributions in disease [61]. To uncover novel associations between cell types and diseases, *deCS* was applied to 51 GWAS data (Table S5) and the association was evaluated by Fisher’s exact test. As shown in **Figure 7**, we observed (1) neuropsychiatric disease, college, and education associated genes being mainly enriched in neurons; (2) immune-related traits enriched in B-cells, T-cells, and NK-cells (lymphoid cells); (3) blood red cell related traits enriched in erythroid cells (Figure S11); (4) bone mineral density enriched in chondrocytes; and (5) high-density lipoproteins enriched in adipocytes. Of note, Alzheimer’s disease, a neurodegenerative disease of the brain, was found to be associated with myeloid lineage cells, rather than neurons. We believe these results collectively shed light on the casual cell type underlying complex traits for both related and unrelated traits. Taken together, the accurate detection of the cell types underlying genetic variants will not only improve our understanding of the molecular mechanisms of complex diseases at the cell type level [62], but also can serve as an important instrumental role in cell type-specific transcriptome-wide association studies [63] and colocalization test [64], among other integrative genetic analyses.

**Figure 7.**
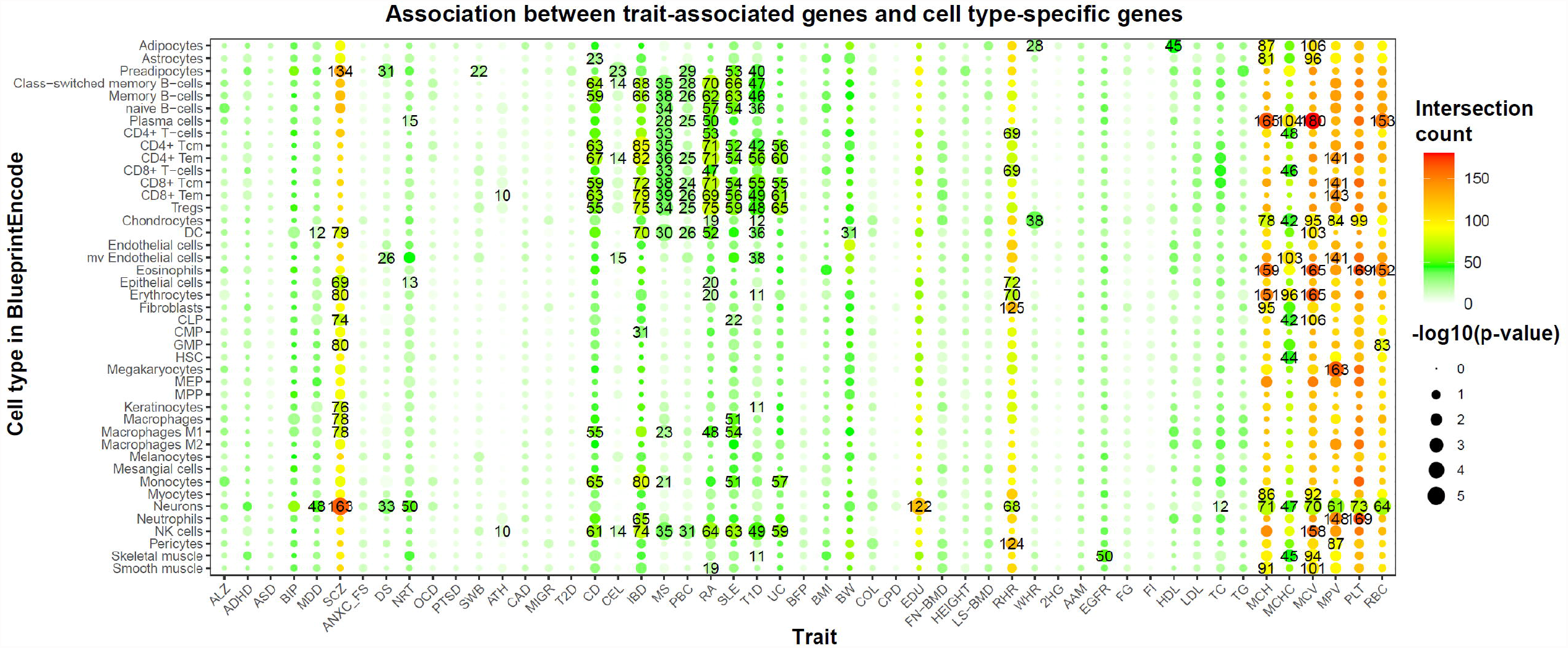
Association between trait-associated genes and cell type-specific genes. Fifty-one traits were analyzed. Heat map shows the significant trait-tissue associations (*p*-value < 0.05). The numbers on the cells indicate the shared gene counts between TAGs, trait-associated genes and CTGenes in BlueprintEncode. The colors and sizes represent the shared gene counts and the -log_10_(transformed *p*-value). Since more significant SNPs are likely found in immune-related diseases and blood-related traits than other traits, we recommend users to focus on the top 3 most relevant cell types for those traits with higher TAGs number.

The second application of *deCS* was applied to bulk RNA expression profiles. Human iPSCs have revolutionized the study of the biological mechanisms underlying psychiatric disorders by establishing cellular models that account for a patient’s genetic background [65]. In most iPSC studies, differentiation quality is routinely assessed [65]. After we normalized the bulk RNA-seq data iPSC-derived neuronal precursor cells (NPCs) and neurons [66], the top matched cell types in 12 of 13 iPSC-NPCs were most enriched in “fetal progenitor cells” or “human embryonic stem cells (hESC)” (Figure S12), while the top matched cell types for iPSC-neurons were most enriched in “fetal neuron”. Therefore, *deCS* can also be applied for decoding the cell type-specificity on bulk RNA-seq.

## Discussion

Single-cell sequencing is a fast-evolving technique that provides deeper insight into the complexity of cellular heterogeneity within the same tissue. One critical step in the analysis is cell type annotation. However, this step is largely done manually, which is time-consuming and subjective. A list of methods was recently developed for automatic cell type annotation [16]. Based on our benchmark results, we demonstrated most classifiers work well when users provide a reference dataset that is conserved with query data. However, due to the small amount of starting tissue material, scRNA-seq data tend to have batch effects [67]. In most condition, researchers do not have enough prior knowledge in the cell type composition of the investigated samples. Thus, finding a suitable reference is time consuming and a subjective task. If some novel cell types were missing in the reference, it would be very difficult to control the sensitivity/specificity of most classifiers, to define them as unknown (novel) cell types, rather than mis-annotate them. To overcome these limitations, we collected multiple human cell type expression profiles from different platforms. As *deCS* only relies on pre-calculated cell-type specificity score and the correlation analysis without further advanced dimension reduction, it can efficiently conduct cell type annotation on each reference respectively. When the cell composition between query and reference datasets is not well conserved (e.g., in the cases of bronchoalveolar, heart, liver and lung in this study), *deCS* achieved the best performance by using the combined highest-scoring labels across multiple reference panels.

*deCS* relies on the predefined cell clusters. There are multiple algorithms for unsupervised clustering, including k-means clustering [68], hierarchical clustering, and graph-based clustering [5], which may generate different clusters and affect *deCS*’s performance. We performed the test across various clustering methods and resolutions, which showed greater than 90% annotation consistency (Table S6). This suggests that *deCS* is robust to predefined cell clusters. In addition, feature selection is also an essential step to obtain the accurate results. In this regard, *deCS* has several advantages over other models. First, *deCS* selects features that co-occur in the reference and query data (discarding useless features in query). Therefore, it drastically reduces the collinearity, the required memory, and runtime for cell annotation, while it achieves comparable accuracy on biologically related cell types (e.g., CD4+, CD8+ T-cells and NK, monocyte, and macrophage). Second, with the expanded references, *deCS* exhibits superior performance to other methods. It is worth noting that even for the same cell type, their expression profiles demonstrated strong heterogeneity among different tissues or different individuals. Thus, it will not work well by simply relying on a single reference. *deCS* can combine the highest-scoring labels across multiple references and dramatically reduce the incorrect cell type annotation when real cell type counterparts are missing in the reference. Although the parameter selection, such as marker genes and correlation measure, may slightly affect the annotation accuracy, the collected comprehensive human cell type profiles in *deCS* provide wider applicability ranging from blood cells to adult and fetal solid tissues. Third, *deCS* supports scRNA-seq cell type annotation for either gene expression profiles or list of genes without prior knowledge. *deCS* is a new tool that it not only benefits the studies of scRNA-seq data, but also yields novel insights for better understanding the molecular basis of thousands of human complex diseases or traits.

We applied *deCS* algorithm to the GWAS data in our recently developed database, Cell type-Specific Enrichment Analysis DataBase (CSEA-DB), which covers 55 unique tissues [69]. Among a total of 10,250,480 trait-cell type associations, we observed significant cell type association in 598 (11.68%) traits. Some human complex diseases or traits were associated with multiple cell types. For example, asthma was found to be associated with both immune and epithelial cells [69]. Moreover, comparing to previous studies [23] that most TAGs enriched in “blood” tissue, *deCS* showed immune TAGs were enriched in lymphoid lineage cells, while red blood cells related traits were enriched in erythroid cells. It has been frequently recognized that pleiotropic effects are ubiquitous in human complex traits [70]. For example, individuals carrying schizophrenia risk alleles tended to be also associated with high risk of Crohn’s Disease [71]. We expect to identify more explicit associations with the increasing high-quality curation of tissue-cell types.

There are several limitations in our study. First, *deCS* does not take tissue information (except for HCAF) into account which may bias the annotation results. For example, 93.2% cells from fetal thymus were annotated as proliferating T-cells (cluster 52), and 85% cells from fetal brain were annotated as fetal neurons (cluster 11) (Table S7). We strongly encourage users to combine prior biological knowledge and apply tissue-matched cell type reference to mitigate potential bias (e.g., almost no neurons or astrocytes in peripheral blood). Second, as a mapping-based annotation toolkit, *deCS* fails to characterize novel cell types. Although we included more than one hundred cell types and most of them come from healthy tissues, it is expected that *deCS* might be challenging for cancer cell annotations due to the expression profile difference between healthy and malignant cells [72]. We believe some “semi-supervised” cell type assignment methods like *scSorter* [73] can address this problem by providing the marker gene in corresponding cell type. Owing to the efforts by the human cell atlas [19] as well as time-dependent change of cell states in gene expression profiles [74], the available large numbers of single cell data sets can be fed in as reference data sets to improve the annotation of future experiments. Despite these limitations, we demonstrated the feasibility and availability of *deCS* for broad applications. Importantly, *deCS* method is not limited to disease gene lists. As our cell type reference panels are only built upon expression profiles, including those unstudied or poorly annotated genes [23, 59], users can highlight the cell type-specificity from any analysis. Taken together, *deCS* is a new tool that not only benefits studies on scRNA-seq data of complex tissues, but also yields novel insights for better understanding the molecular basis of various human diseases and traits.

## Code availability

The source codes and results are implemented in an R package and are freely available at GitHub: https://github.com/bsml320/deCS, or BioCode at NGDC with Accession <u>BT007286</u>. We also developed a shiny application as part of the deCS package, and it is available at https://gpei.shinyapps.io/decs_cor/ and https://gpei.shinyapps.io/decs_fisher/.

## CRediT author statement

**Guangsheng Pei**: Conceptualization, Methodology, Software, Resources, Data Curation, Visualization, Writing - original draft. **Fangfang Yan**: Data Curation, Visualization, Validation, Writing - review & editing. **Lukas M. Simon**: Validation, Writing - review & editing. **Yulin Dai**: Validation, Writing - review & editing. **Peilin Jia**: Conceptualization, Methodology, Writing - review & editing. **Zhongming Zhao**: Conceptualization, Methodology, Project administration, Supervision, Writing - review & editing, Funding acquisition. All authors read and approved the final manuscript.

## Competing interests

The authors declare no competing interests.

## Acknowledgments

The authors thank all the members the Bioinformatics and Systems Medicine Laboratory (BSML). We also thank those investigators who generated the reference data. This study was partially supported by National Institutes of Health grants (R01LM012806, R01DE030122, and R01DE029818). We thank the resource support from Cancer Prevention and Research Institute of Texas (CPRIT RP180734 and RP210045). Funding for open access charge: CPRIT (RP180734).

## Supplementary material

**Figure S1 Summary of three bulk-RNA derived cell type references**

**A-C**. *Spearman’s rank* correlation analysis for each cell type by using the average expression for each gene. **D-F**. Uniform Manifold Approximation and Projection (UMAP) dimension reduction analysis investigated the global landscape of cell type expression profiles. **G-I**. Variance comparison of different gene features. Each dot represents one gene feature. *X*-axis: type of gene feature (e.g. protein coding, lincRNA). *Y*-axis: the variance of gene feature, scaled by absolute abundance.

**Figure S2 Shared cell type-specific genes comparison between bulk-RNA and scRNA-seq derived references**

**A-C**. Main CTGenes comparison between BlueprintEncode and ImmuneCellExpression, BlueprintEncode and MonacoImmune, ImmuneCellExpression and MonacoImmune. **D-F**. Fine CTGenes comparison between BlueprintEncode and ImmuneCellExpression, BlueprintEncode and MonacoImmune, ImmuneCellExpression and MonacoImmune. **G**. CTGenes comparison between BlueprintEncode and human cell landscape. The number indicates the shared CTGenes number between two cell types.

**Figure S3 Overview of *deCS* flowchart for scRNA-seq annotation**

We recommend users apply *Seurat* standard pre-processing workflow, including transcription noise cells cleaning, highly variable genes identification, dimension reduction and cell clustering analysis. Then, *Seurat* FindAllMarkers() function can use to identify cluster-specific marker genes, and AverageExpression() function can use to calculate the gene average expression level for each cell cluster. Base on provided *t*-statistics and *z*-score for each gene, we implemented two test approaches for scRNA-seq cell type annotation. Strategy A: user can define genes with the highest *t*-statistics or *z*-score (top 5%) as CTGenes, then investigate whether a set of “cluster-specific marker genes” of interest are disproportionately overlapped with CTGenes by using Fisher’s exact test. Strategy B: user can calculate the *Pearson correlation coefficient* (PCC) or *Spearman’s rank correlation coefficient* between query scaled expression profiles and *t*-statistics (or *z*-score) of each cell type in reference, then assigned the label with the highest score to query profile.

**Figure S4 Cell type annotation of bronchoalveolar immune cells data set by using different references**

**A**. BlueprintEncode. **B**. Human cell landscape. **C**. MonacoImmune. *X*-axis: 10 cell type clusters (query). *Y*-axis: the reference cell types. The colors representing the PCC or genes intersection ratio, and the sizes representing the -log_10_ transformed *p*-value. Non-significant associations (*p*-value > 0.05) were replaced by blank color.

**Figure S5 Cell type annotation of cardiac cells from human fetal cardiac cells by using different reference or correlation measurement**

**A-C**. *Pearson correlation coefficient* on BlueprintEncode, human cell landscape and human cell atlas of fetal. **D-F**. *Spearman’s correlation coefficient* on BlueprintEncode, human cell landscape and human cell atlas of fetal. *X-*axis: 9 cell type clusters (query). *Y*-axis: the reference cell types. The colors representing the PCC or *Spearman’s correlation coefficient*, and the sizes representing the -log_10_ transformed *p*-value. Non-significant associations (*p*-value > 0.05) were replaced by blank color.

**Figure S6 Overview of cell type trajectory in human immune system**

**Figure S7 Hierarchical clustering analysis of different cell type in each reference**.

Numbers at the nodes indicate bootstrap values for 100 replicates.

**Figure S8 Violin plot shows the expression probability distributions across each cluster’s canonical marker genes (top 1) in PBMC data set**

**Figure S9 Single cell annotation results by CHETAH model**

**A**. Top 30 marker genes for each cell cluster. **B**. Top 2000 most variable genes. **C**. All detected genes.

**Figure S10 Principal component analysis (PCA) of down-sampled bulk RNA-seq data set**

**A**. Sequencing depth identity. **B**. Sample identity based on raw TPM value. Z-score normalization by **C**. sequencing depth and **D**. sample identity. S1 to S5 denote five samples NSC353, NSC412, NSC413, NSC416 and NSC419, respectively.

**Figure S11 Association between 51 trait-associated genes and cell type-specific genes**

Heat map shows the significant trait-tissue associations (*p*-value < 0.05). The numbers on the cells indicate the shared gene counts between trait-associated genes and CTGenes in human cell landscape. The colors and sizes represent the shared gene counts and the -log_10_(transformed *p*-value). Since more significant SNPs are likely found in immune-related diseases and blood-related traits than other traits, we recommend users to focus on the top 3 most relevant cell types for those traits with higher TAGs number.

**Figure S12 *deCS* annotation on bulk RNA-seq of iPSC-derived cell lines**

iPSC-NPCs (n = 13) and iPSC-neurons (n = 11).

**Table S1 Gene Ontology (GO) and KEGG pathway enrichment analysis of cell type-specific genes**

**Table S2 *deCS* annotation results on 6 different solid tissues**

**Table S3 Performance comparison of *deCS* and other software on PBMC data set**

**Table S4 Simulated TPM expression profiles of 5 down-sampled bulk RNA-seq samples**

**Table S5 GWAS summary and statistics of the genes associated with regulatory loci in 51 human complex traits**

**Table S6 Consistency comparison of *deCS* annotation results between different clustering methods or under different clustering resolutions**

**Table S7 Summary of cell type composition of major human organs from human cell landscape reference**

## Notes

### Competing Interest Statement

The authors have declared no competing interest.

### Summary of Updates

1. We further benchmarked two methods from Abdelaal et al. study and the performance (running time and accuracy rate) of deCS was still impressive on either PBMC or various tissue data sets. 2. To increase its applicability, so we further incorporated a shiny application (at https://gpei.shinyapps.io/decs_cor/ or https://gpei.shinyapps.io/decs_fisher/) to deCS package. 3. We added more discussion on the strengths and weaknesses of deCS, and also reorganized Figure 1 to better describe the detailed steps to pre-process the data. 4. English writing and format of the manuscript have been enhanced by native speakers.

