## Supplementary figures and images for "*deCS*: A Tool for Systematic Cell Type Annotations of Single-cell RNA Sequencing Data among Human Tissues"

### Figure S1

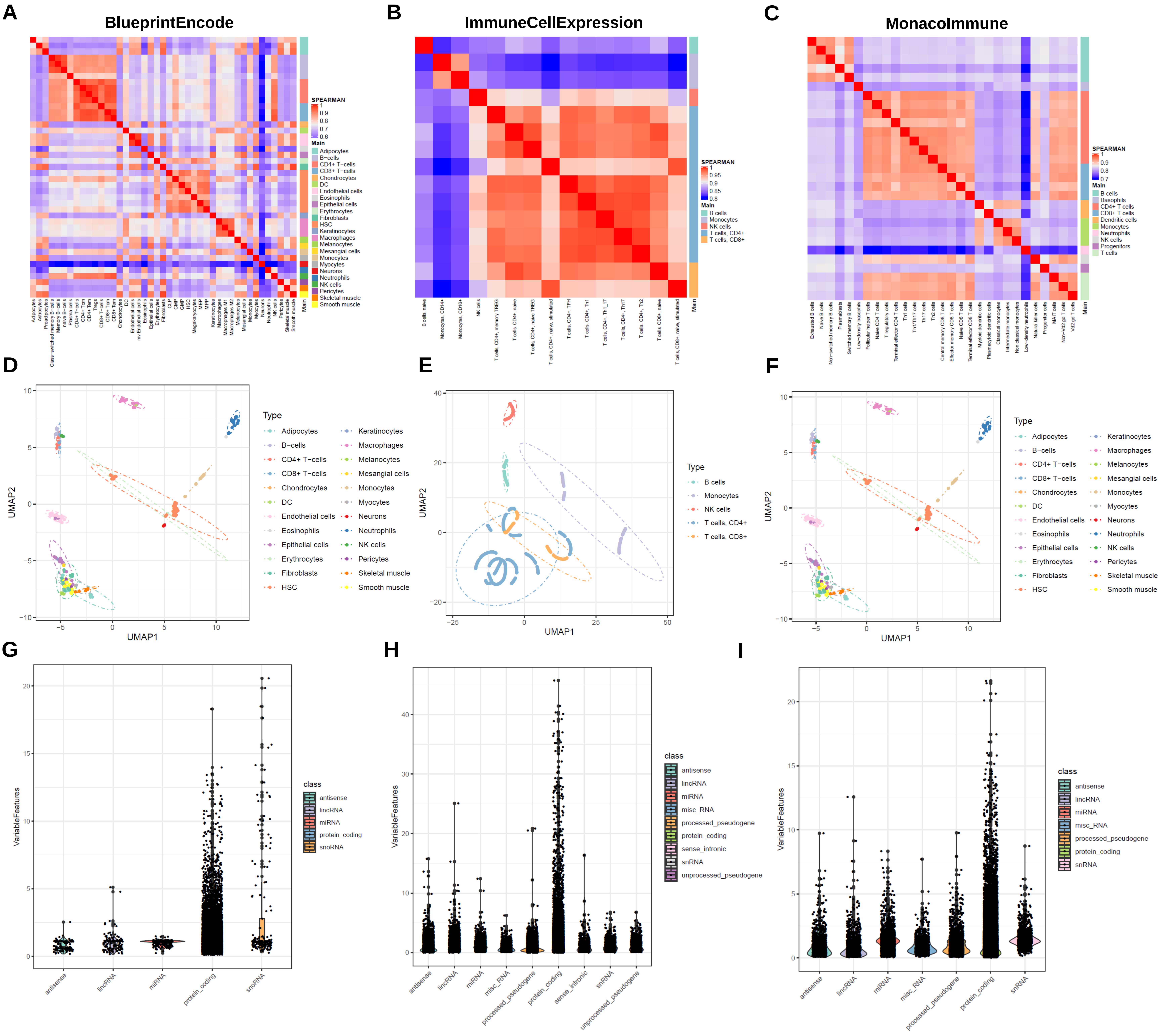

### Figure S2

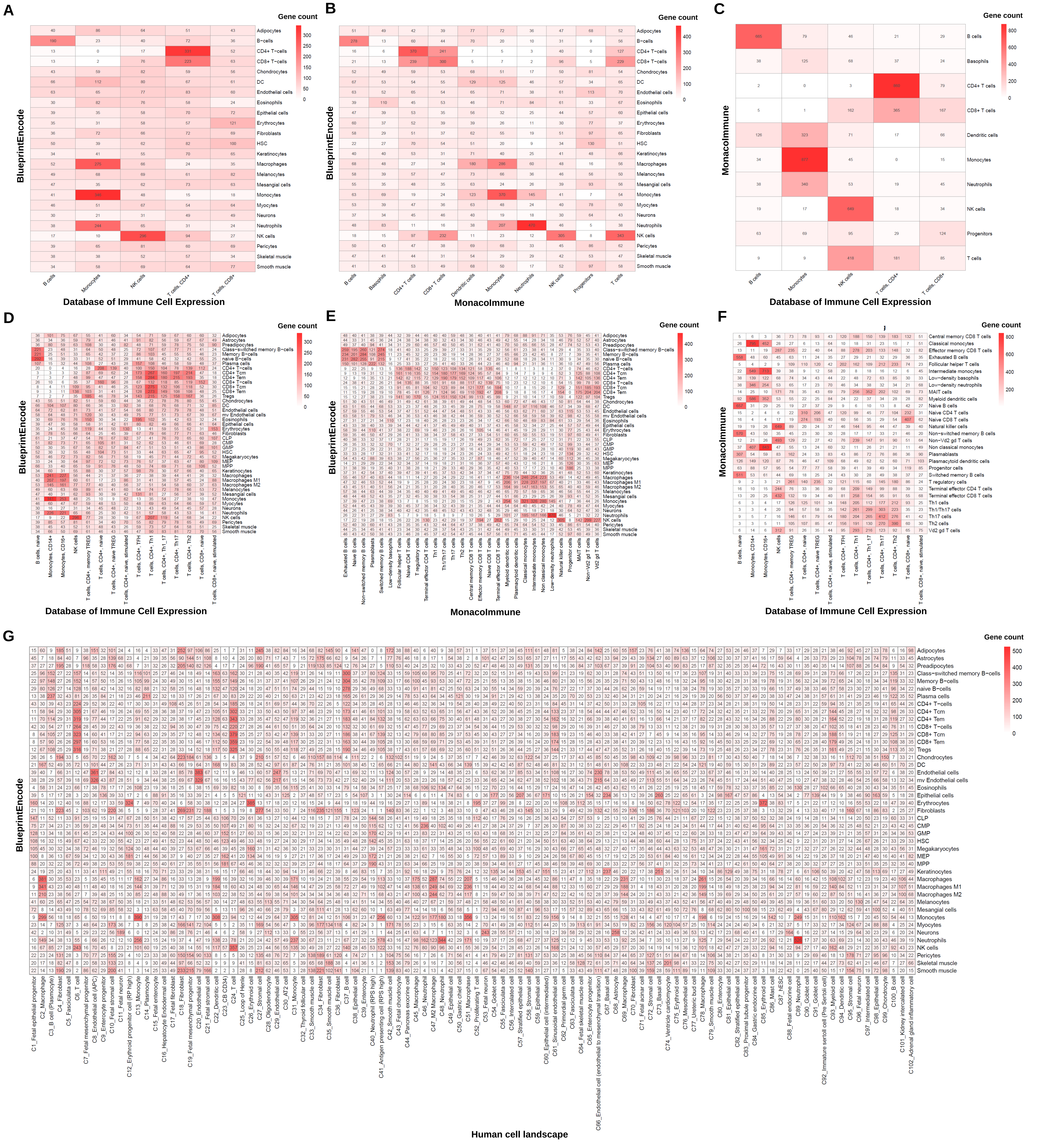

### Figure S3

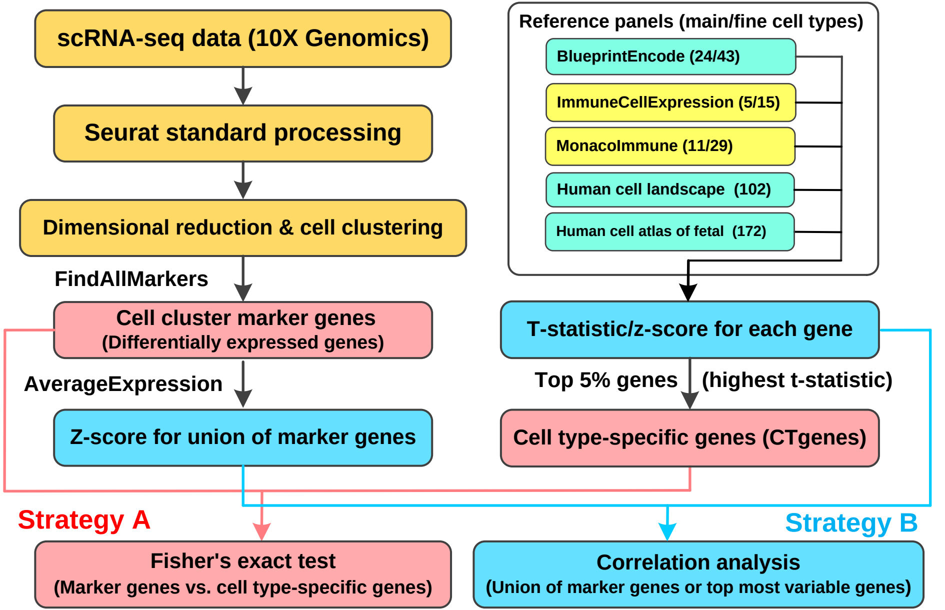

### Figure S4

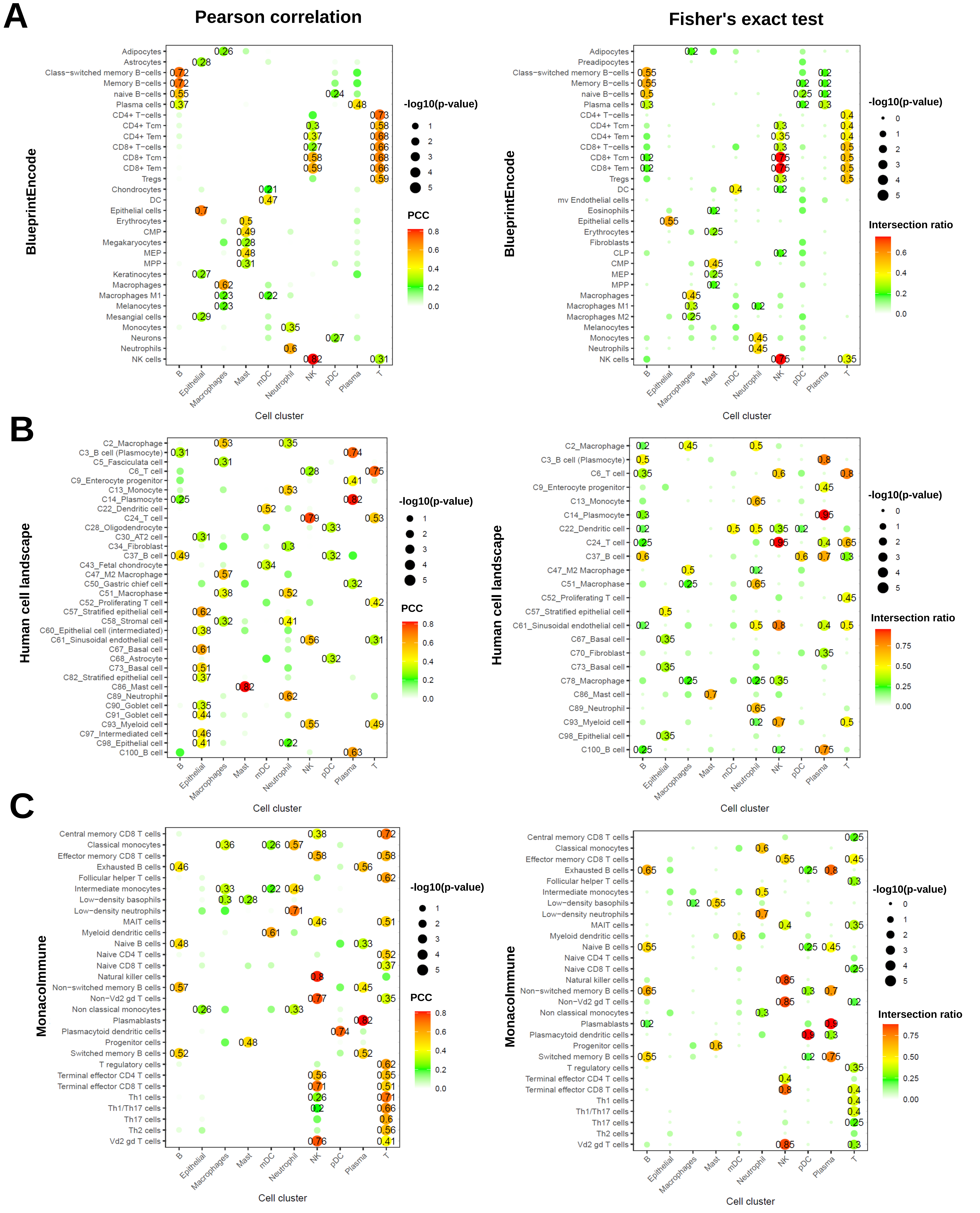

### Figure S5

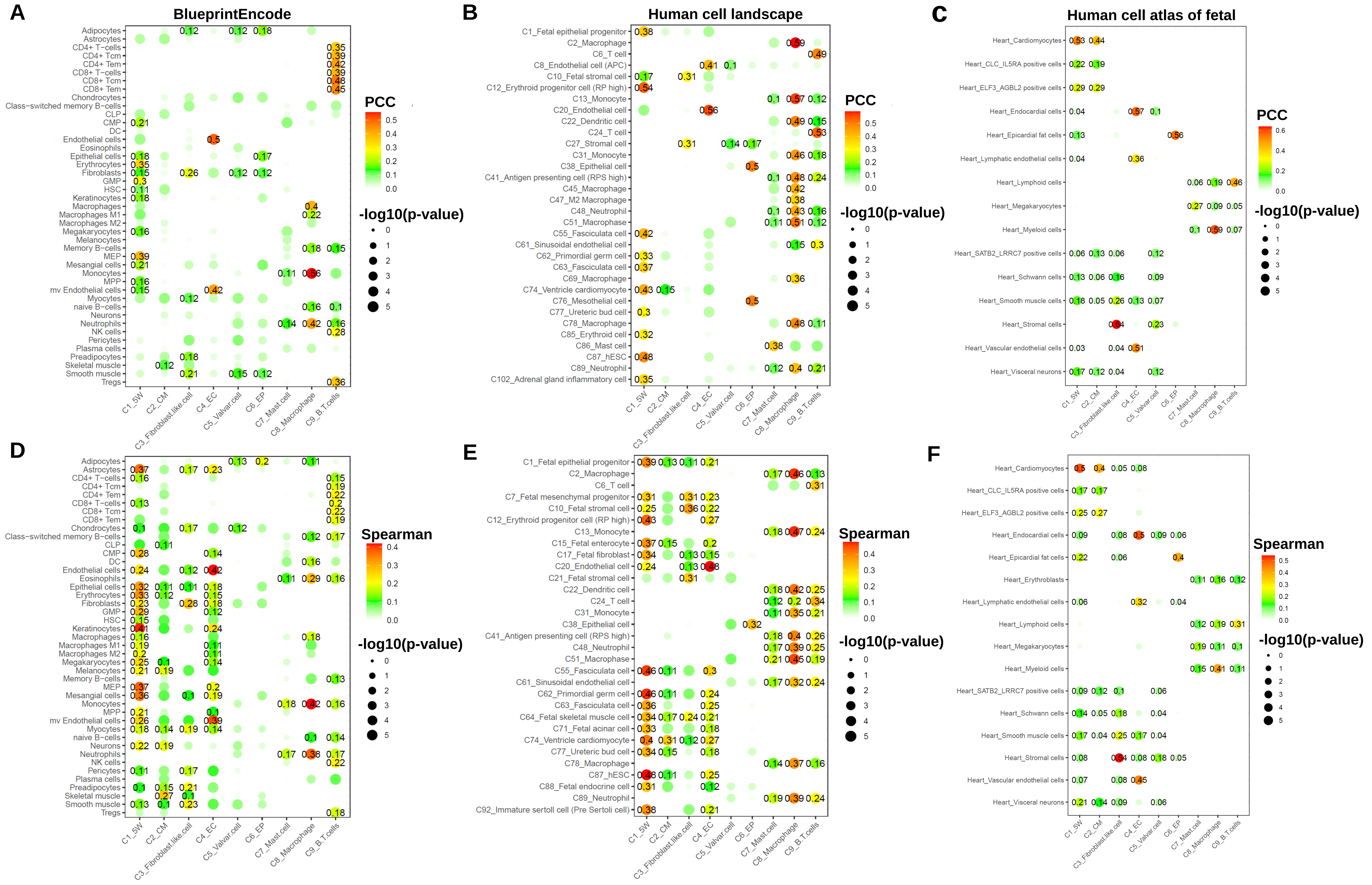

### Figure S6

#### Overview of immune system cell types

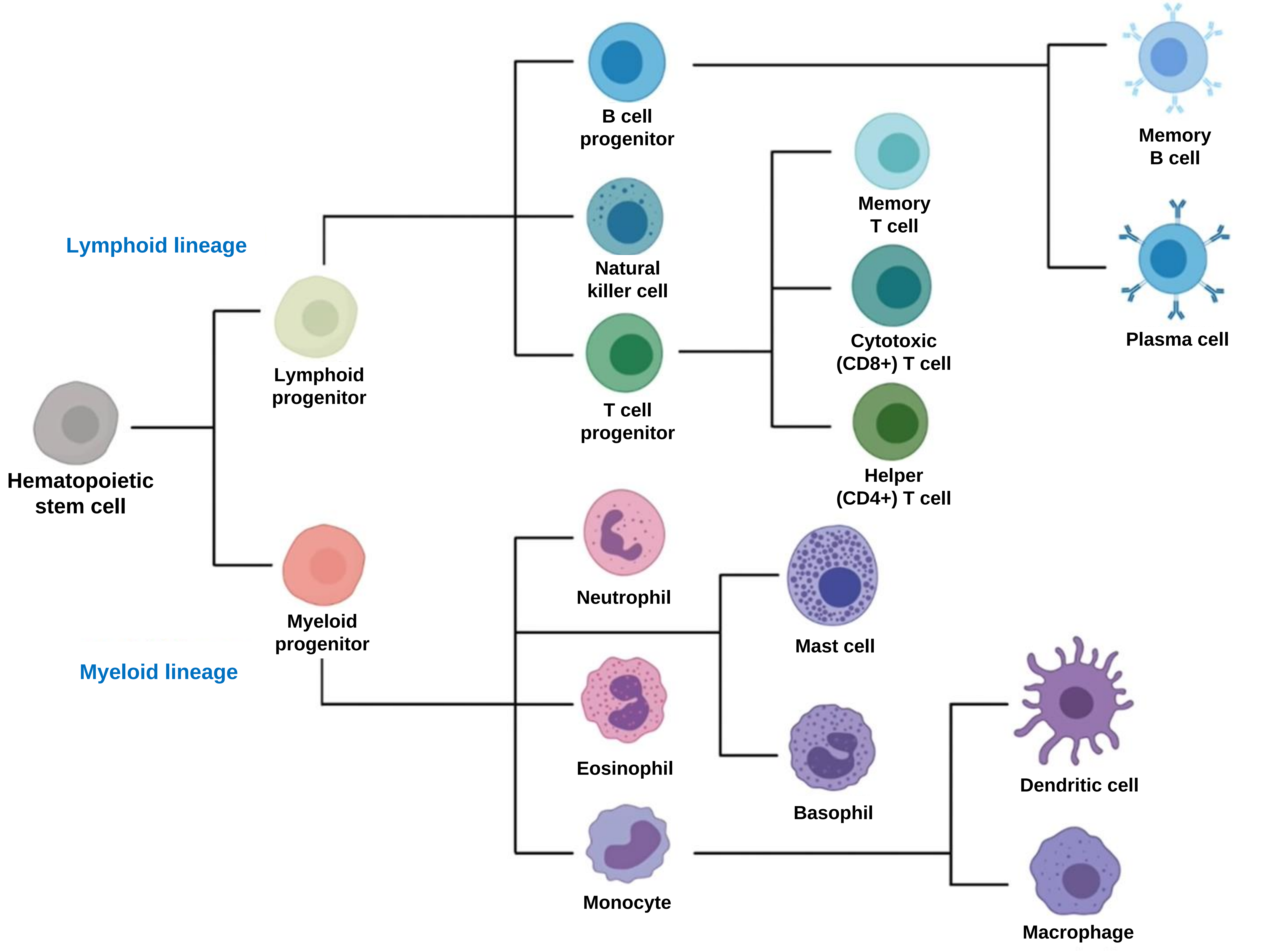

### Figure S7

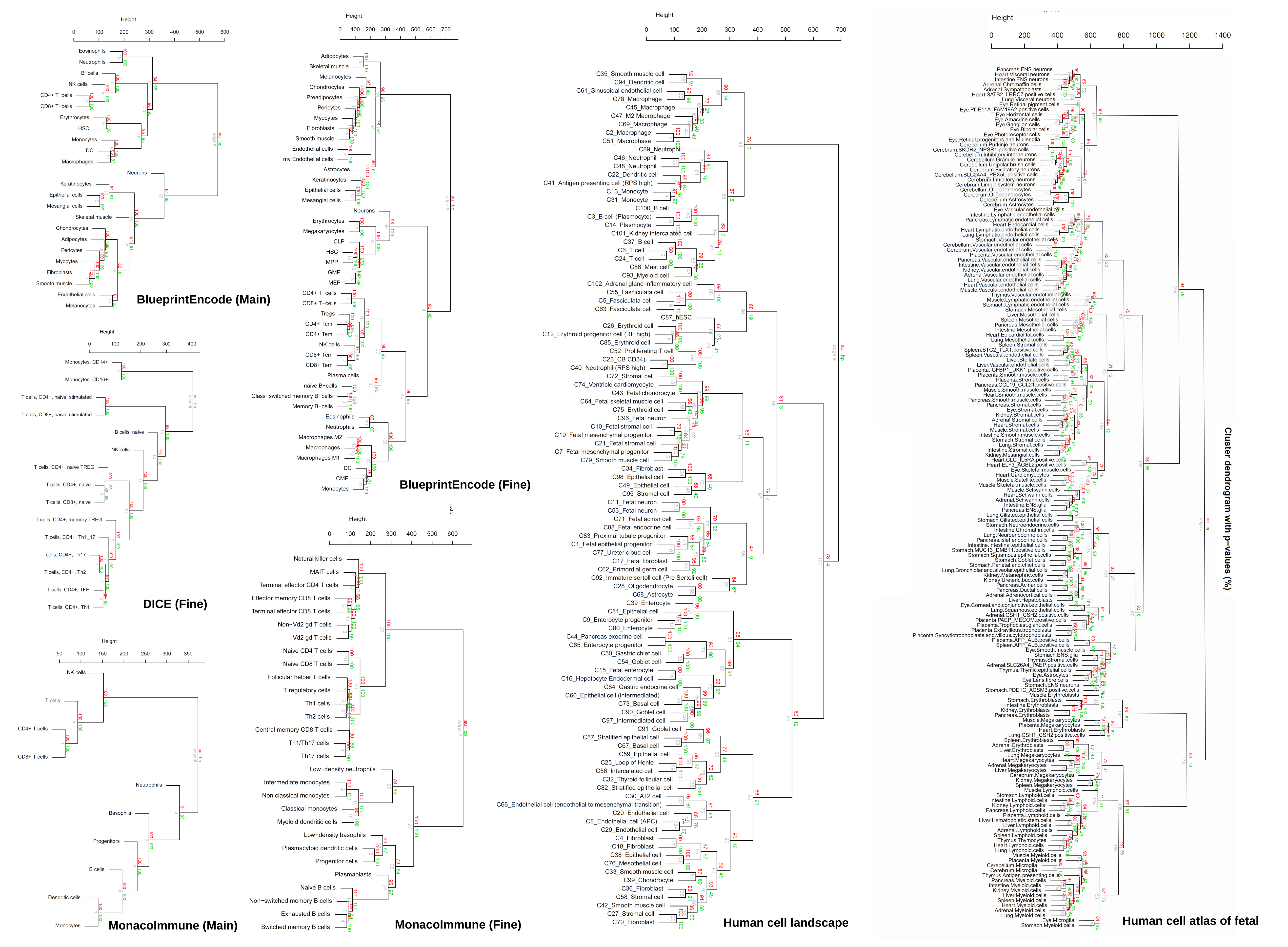

### Figure S8

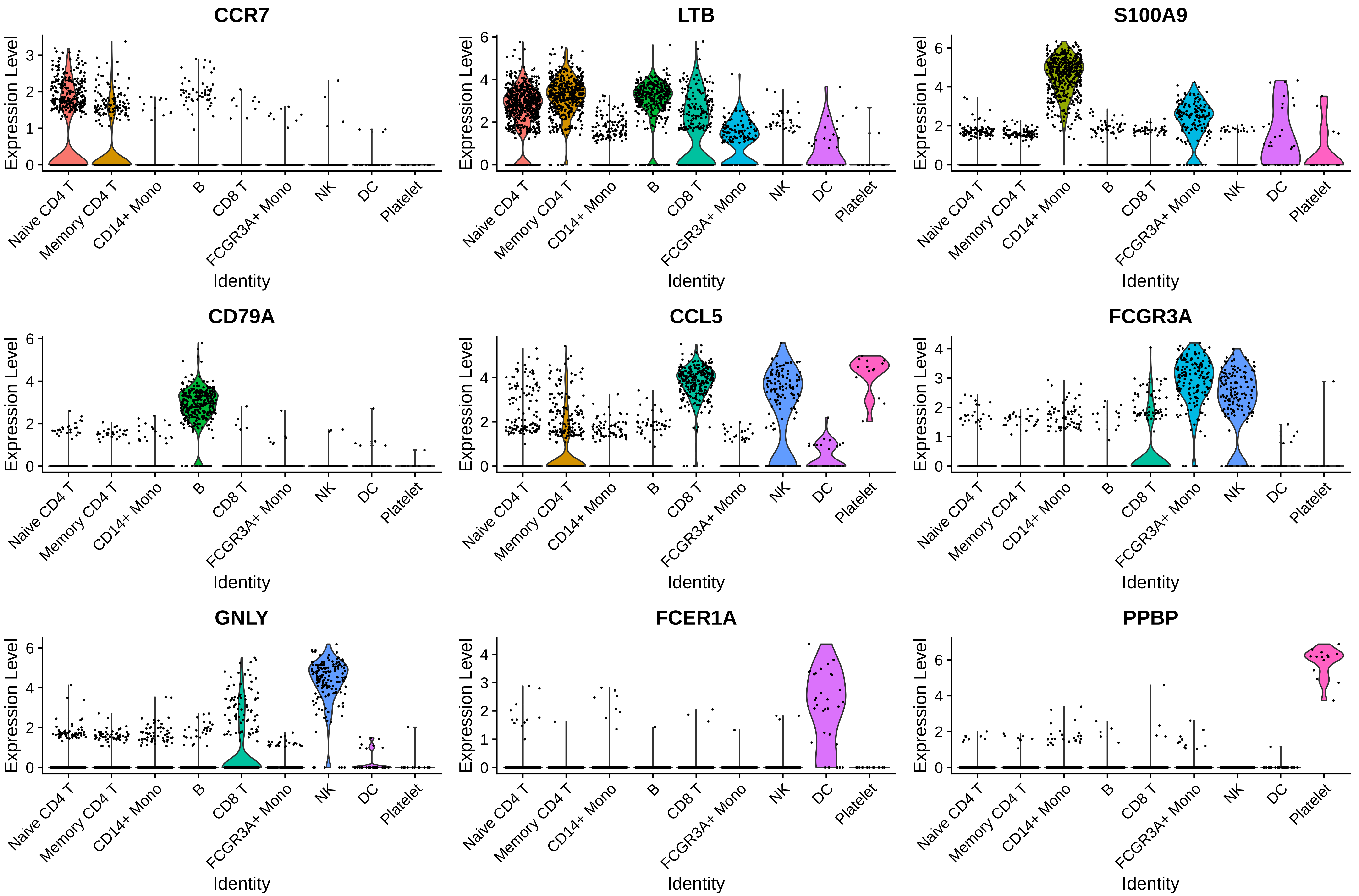

### Figure S9

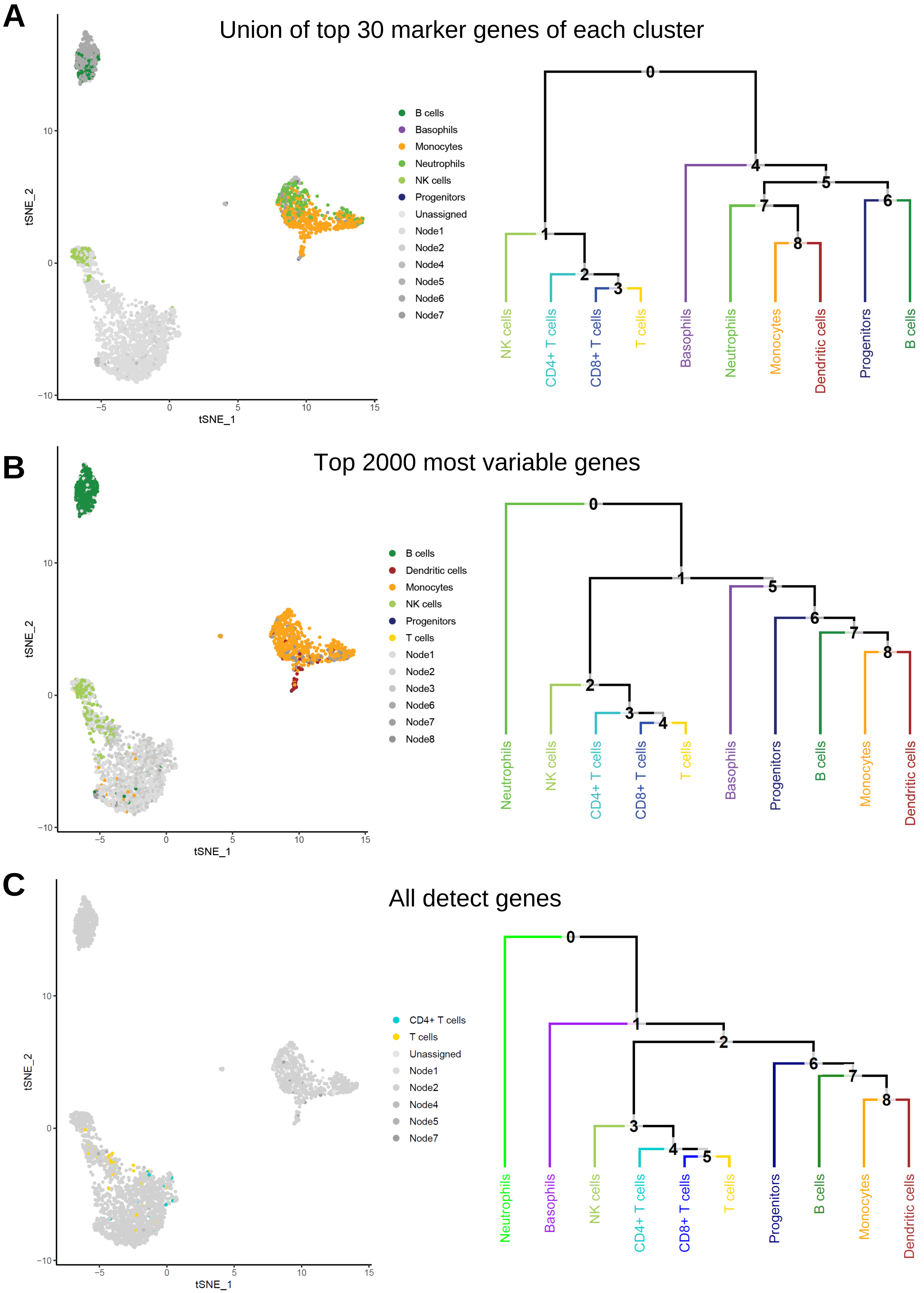

### Figure S10

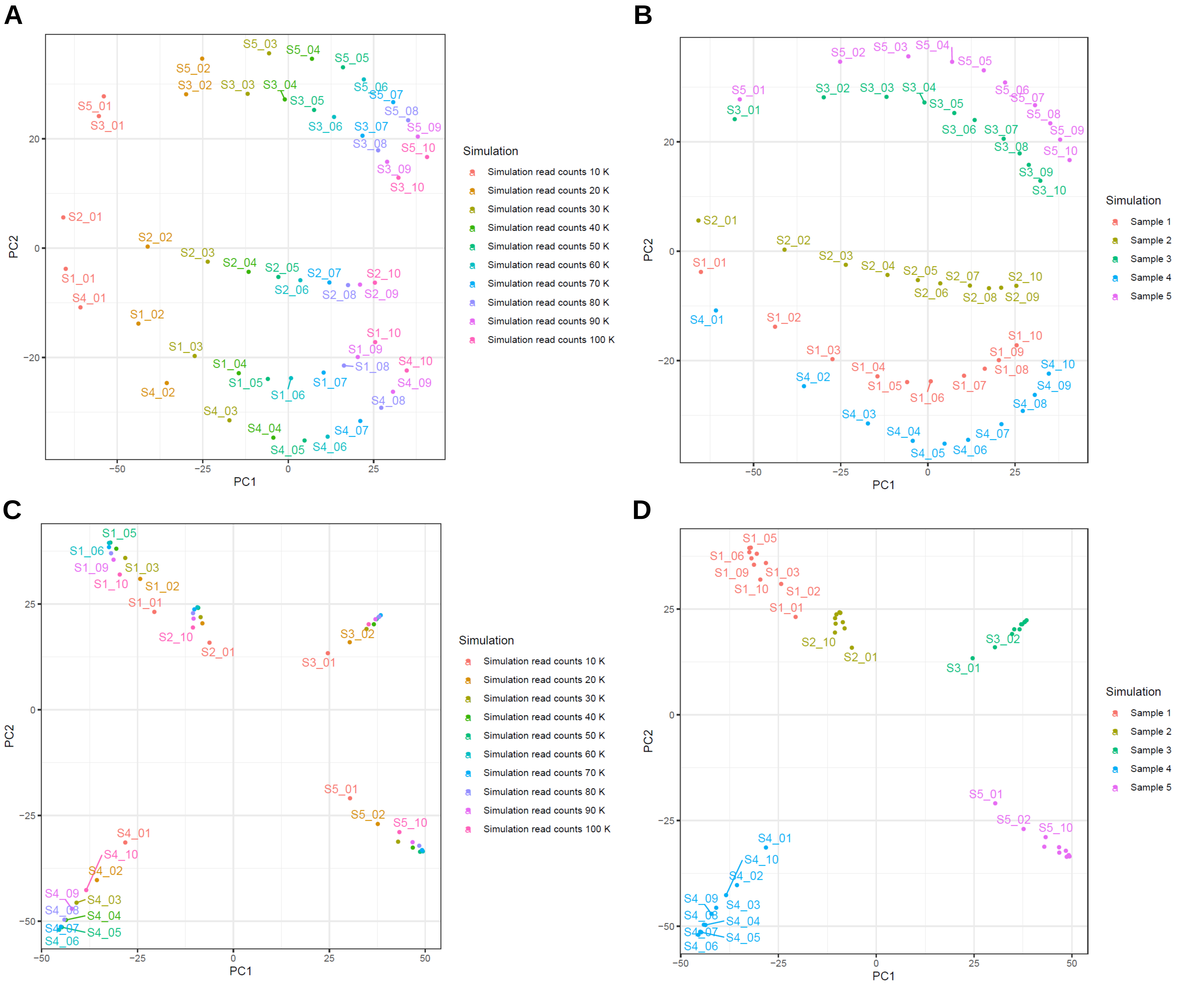

### Figure S11

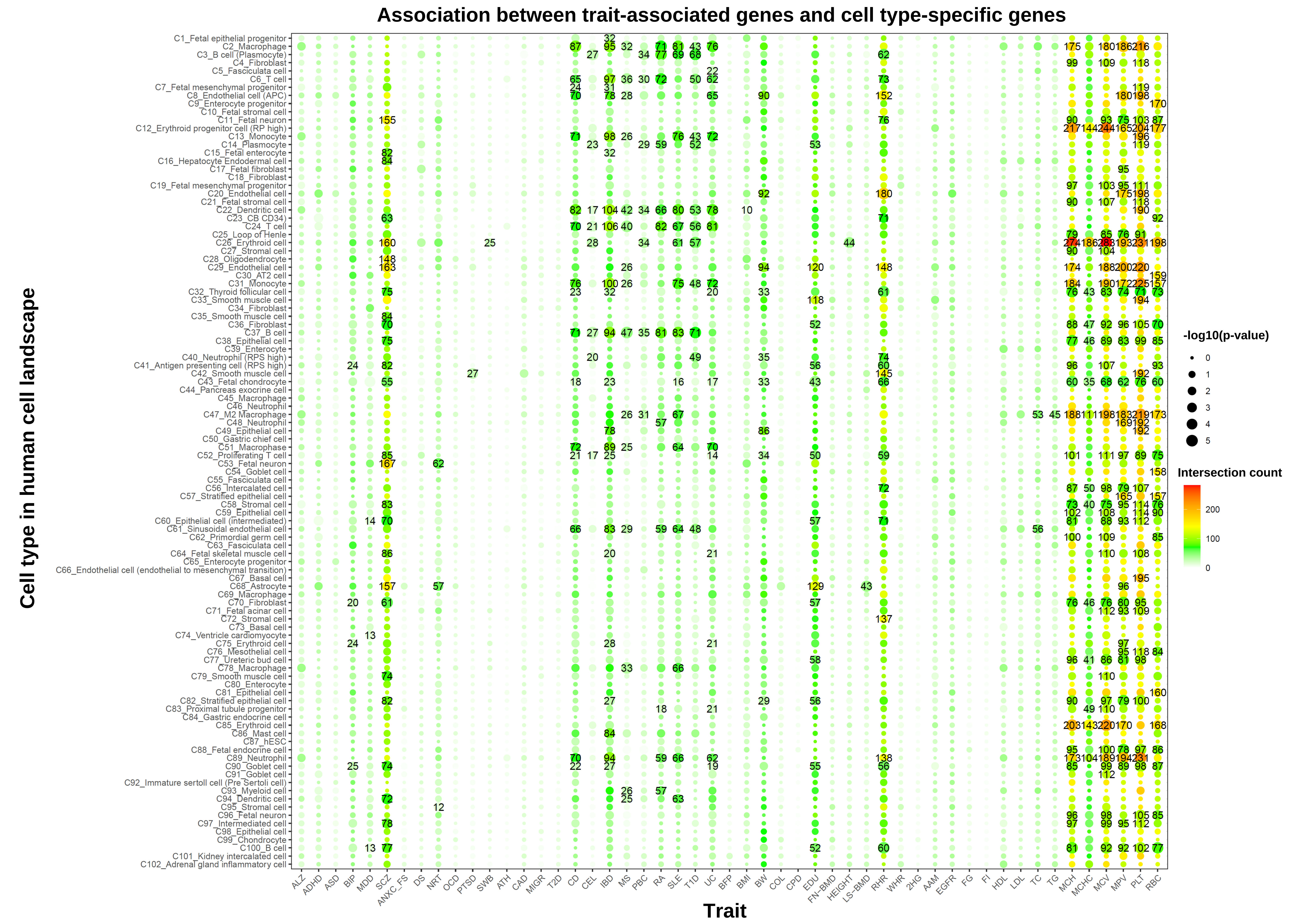

### Figure S12

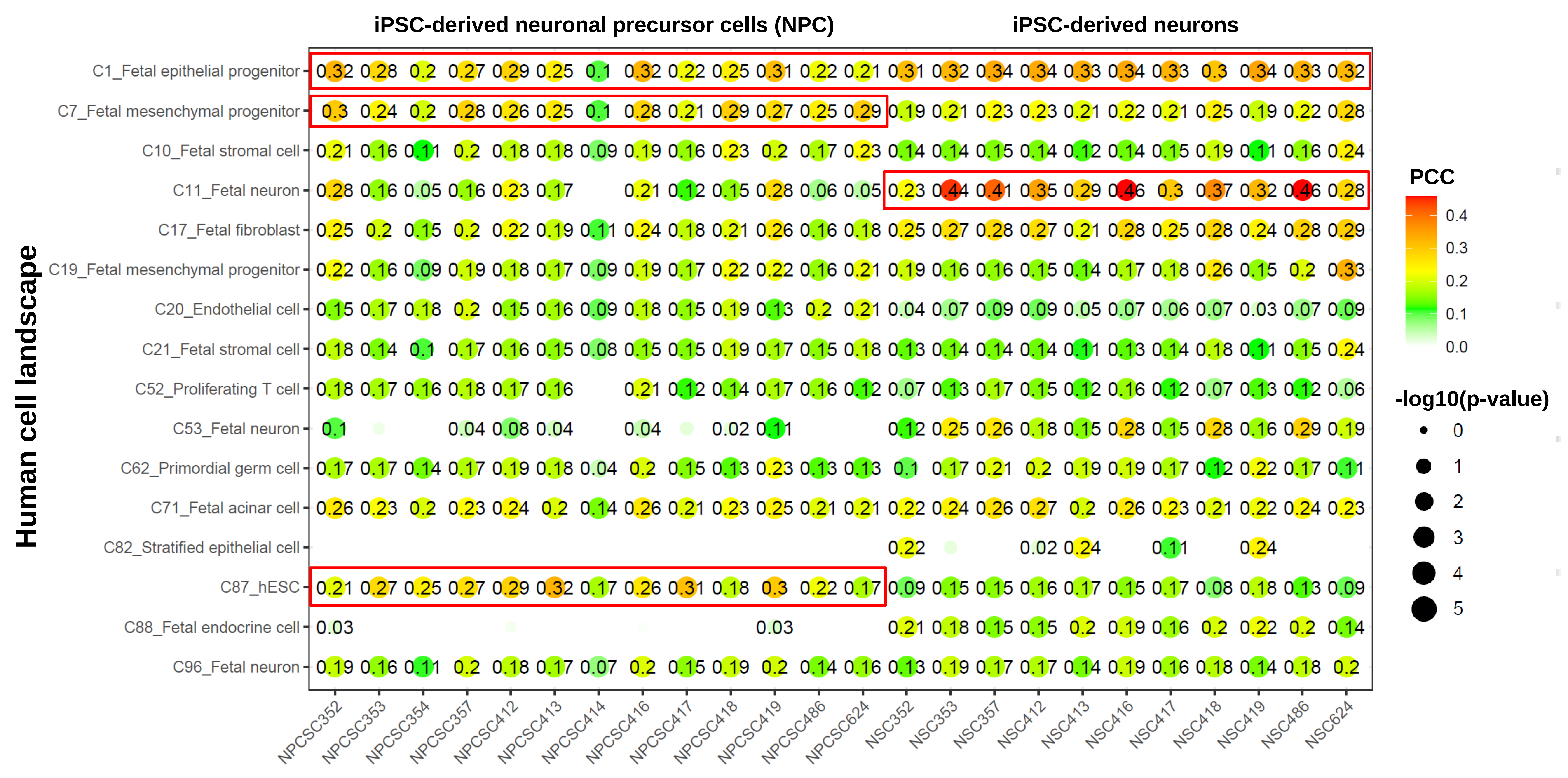
